# ARHGAP12 suppresses F-actin assembly to control epithelial tight junction mechanics and paracellular leak pathway permeability

**DOI:** 10.1101/2024.08.24.609485

**Authors:** Hana Maldivita Tambrin, Yun Liu, Kexin Zhu, Xiang Teng, Yusuke Toyama, Yansong Miao, Alexander Ludwig

**Affiliations:** School of Biological Sciences, Nanyang Technological University Singapore, 60 Nanyang Drive, Singapore 637551; NTU Institute of Structural Biology, Nanyang Technological University Singapore, 59 Nanyang Drive, Singapore 636921; Mechanobiology Institute, National University of Singapore, 5A Engineering Drive, Singapore 117411

## Abstract

Tight junctions (TJ) separate body compartments and control the paracellular transport of ions, solutes, and macromolecules across epithelial barriers. There is evidence that claudin-based ion transport (the pore pathway) and the paracellular transport of macromolecules (the leak pathway) are independently regulated processes. However, how leak pathway permeability is controlled is not well understood. Here we have identified the Cdc42/Rac GTPase activating protein ARHGAP12 as a novel and specific regulator of the leak pathway. ARHGAP12 is recruited to TJ via an interaction between its SH3 domain and the TJ protein ZO-2. Using a combination of biochemical and biophysical approaches, in vitro actin polymerisation assays, and permeability assays in MDCK-II cells, we show that ARHGAP12 suppresses N-WASP-mediated F-actin assembly at TJ to dampen junctional tension. This promotes paracellular leak pathway permeability without affecting ion flux. Mechanistically, we demonstrate that the ARHGAP12 tandem WW domain interacts directly and in a multivalent manner with an array of PPxR motifs in the proline-rich domain of N-WASP. This interaction is sufficient to suppress SH3 domain-mediated N-WASP oligomerisation and Arp2/3-driven F-actin assembly in vitro. Collectively our data demonstrate a critical role for ARHGAP12 in suppressing junctional F-actin assembly and tension to promote the flux of small macromolecules across the TJ.

## Introduction

Tight junctions (TJ) are critical for animal development and physiology. They seal the paracellular space between adjacent epithelial cells and thereby form a barrier that physically separates body compartments. At the same time, the TJ acts as a gate that selectively controls the paracellular flux of ions, water and various macromolecules across epithelial tissues. The precise control of the permeability barrier is critical for tissue homeostasis and organ function, defects of which can cause a range of diseases including intestinal bowel disease and hypomagnesemia (Horowitz et al., 2023; Hou, 2014; Krug et al., 2014).

TJ are composed of integral membrane proteins that form a complex network of intramembranous fibrils, the so-called TJ strands. TJ strands are formed by the claudin family, occludin, and other integral membrane proteins, which interact via their C-terminal tails with cytoplasmic scaffolding proteins, such as the Zonula Occludens (ZO) family, cingulin and paracingulin. ZO and cingulin proteins link the TJ to the F-actin cytoskeleton, control TJ formation, dynamics, and mechanics, and mediate signaling in response to mechanical cues (Citi, 2019; Matter and Balda, 2003; Otani and Furuse, 2020; Zihni et al., 2016). Recent data indicate that TJ are under mechanical load and sense and transmit mechanical forces (Haas et al., 2020; Rouaud et al., 2023; Spadaro et al., 2017). Such forces are generated by the junction-associated actin cytoskeleton, which is regulated by the RhoGTPases Rho, Rac, and Cdc42 (Bock et al., 2021; Diaz-Diaz et al., 2020; Elias et al., 2015; Hodge and Ridley, 2016; Pichaud et al., 2019). The activities of RhoGTPases are tightly controlled by a large number of Rho Guanine Exchange Factors (RhoGEFs) and Rho GTPase Activating Proteins (RhoGAPs) (Citi et al., 2011; Hodge and Ridley, 2016; Ngok et al., 2014). Several RhoGEFs have been implicated in the control of TJ formation (Chen and Macara, 2005; Mertens et al., 2005) or maturation (Zihni et al., 2014), as well as in the regulation of junctional actomyosin contractility (Benais-Pont et al., 2003; Guillemot et al., 2008; Haas et al., 2020; Itoh et al., 2012; Nakajima and Tanoue, 2011; Scott et al., 2016; Terry et al., 2011). However, barring few exceptions (Guillemot et al., 2014; Wells et al., 2006), we know little about the nature and functions of RhoGAPs at TJ.

TJ control paracellular flux via two distinct pathways, the pore and the leak pathway (Monaco et al., 2021; Shen et al., 2011). Whilst it is well established that the claudin family mediates the paracellular transport of specific ions and small uncharged molecules via the pore pathway (Colegio et al., 2002; Tsukita et al., 2019), the molecular nature and regulation of the leak pathway are less clear. The leak pathway is a low capacity, non-charge selective route for molecules up to 10 nm in diameter. There is some evidence that such macromolecules are specifically transported through tricellular junctions (Cho et al., 2022; Isasti-Sanchez et al., 2021). Another prominent model posits that the leak pathway is mediated by transient breaks in the TJ strand barrier. Such breaks could be elicited through mechanical forces acting on the TJ, changes in TJ composition or remodelling of the TJ structure, or a combination of these factors (Monaco et al., 2021; Shen et al., 2011; Varadarajan et al., 2019). For instance, cell stretching (Cavanaugh et al., 2006; Cohen et al., 2008; Samak et al., 2014) or activation of myosin light chain kinase (He et al., 2020; Horowitz et al., 2023; Shen et al., 2006; Turner et al., 1997) promotes the leak pathway by increasing junctional tension. Inactivation of ZO-1 or both ZO-1 and ZO-2 also causes a marked increase in junctional contractility and elevates the permeability to small macromolecules (Belardi et al., 2020; Choi et al., 2016; Spadaro et al., 2017; Van Itallie et al., 2009). This indicates that ZO proteins support the TJ barrier by suppressing junctional actomyosin activity. However, increases in leak pathway permeability have been observed under conditions that both increase and decrease junctional tension (Monaco et al., 2021). For instance, loss or inhibition of the actin nucleation promoting factor (NPF) N-WASP, the Cdc42 effector TOCA-1, or Arp2/3 complex (which nucleates branched actin networks downstream of N-WASP/TOCA-1) decreases junctional tension and increases flux through the TJ leak pathway (Garber et al., 2018; Van Itallie et al., 2015; Wu et al., 2014). Taken together the data suggest that the junctional F-actin cytoskeleton and actomyosin-mediated tension need to be finely balanced to permit the flux of macromolecules across the TJ without compromising the TJ barrier. However, to date there is little mechanistic insight into how this is achieved.

Here we show that the Rac/Cdc42 GAP ARHGAP12 controls junctional tension and TJ leak pathway permeability by suppressing N-WASP-mediated branched actin polymerisation via Arp2/3 complex. ARHGAP12 is recruited to TJ through an interaction between its SH3 domain and ZO-2. The tandem WW domain of ARHGAP12 interacts directly and in a multivalent manner with the proline-rich domain (PRD) of N-WASP, and this interaction alone is sufficient to block N-WASP activity and Arp2/3-mediated F-actin assembly in vitro. CRISPR/Cas9-mediated inactivation of *ARHGAP12* in MDCK-II cells delays TJ formation, increases junctional tension, and reduces the flux of macromolecules across mature epithelial monolayers. Taken together our data indicate that ARHGAP12 plays an important role in suppressing F-actin assembly and tension at TJ to promote paracellular transport through the TJ leak pathway.

## Results

### ARHGAP12 is recruited to tight junctions via ZO-2

Using proximity proteomics we previously identified ARHGAP12 as a component of TJ in MDCK-II cells (Tan et al., 2020) (Figure S1). Two isoforms of ARHGAP12 have been reported, ARHGAP12a and ARHGAP12b (Zhang et al., 2002). Both isoforms contain an N-terminal Src homology (SH3) domain, a tandem WW (tWW) domain, a pleckstrin homology (PH) domain, and a C-terminal GAP domain. ARHGAP12 specifically suppresses Rac and Cdc42 (Ba et al., 2016; Diring et al., 2019; Gentile et al., 2008; Schlam et al., 2015) and has been reported to localise to the apical junctional complex in various epithelial tissues (Matsuda et al., 2008). However, the functions of this protein in epithelial cells are unknown.

To ascertain that ARHGAP12 localizes to TJ, MDCK-II cells grown on Transwell filters were fixed and stained with antibodies against ARHGAP12 and Pals1 (an apical junctional marker). X/y and orthogonal projections of MDCK-II monolayers indicated partial co-localization of ARHGAP12 with Pals1 (Figure 1A and 1B). GFP fusion proteins of ARHGAP12a (and ARHGAP12b, data not shown) as well as a construct lacking the C-terminal PH and GAP domains (GFP-SH3-tWW) stably expressed in MDCK-II cells also colocalised with the TJ markers Par3 and ZO-1 (Figure 1C and 1D). We concluded that both isoforms of ARHGAP12 are localised to TJ and that the N-terminal SH3-tWW module is sufficient for TJ localisation.

**Figure 1:**
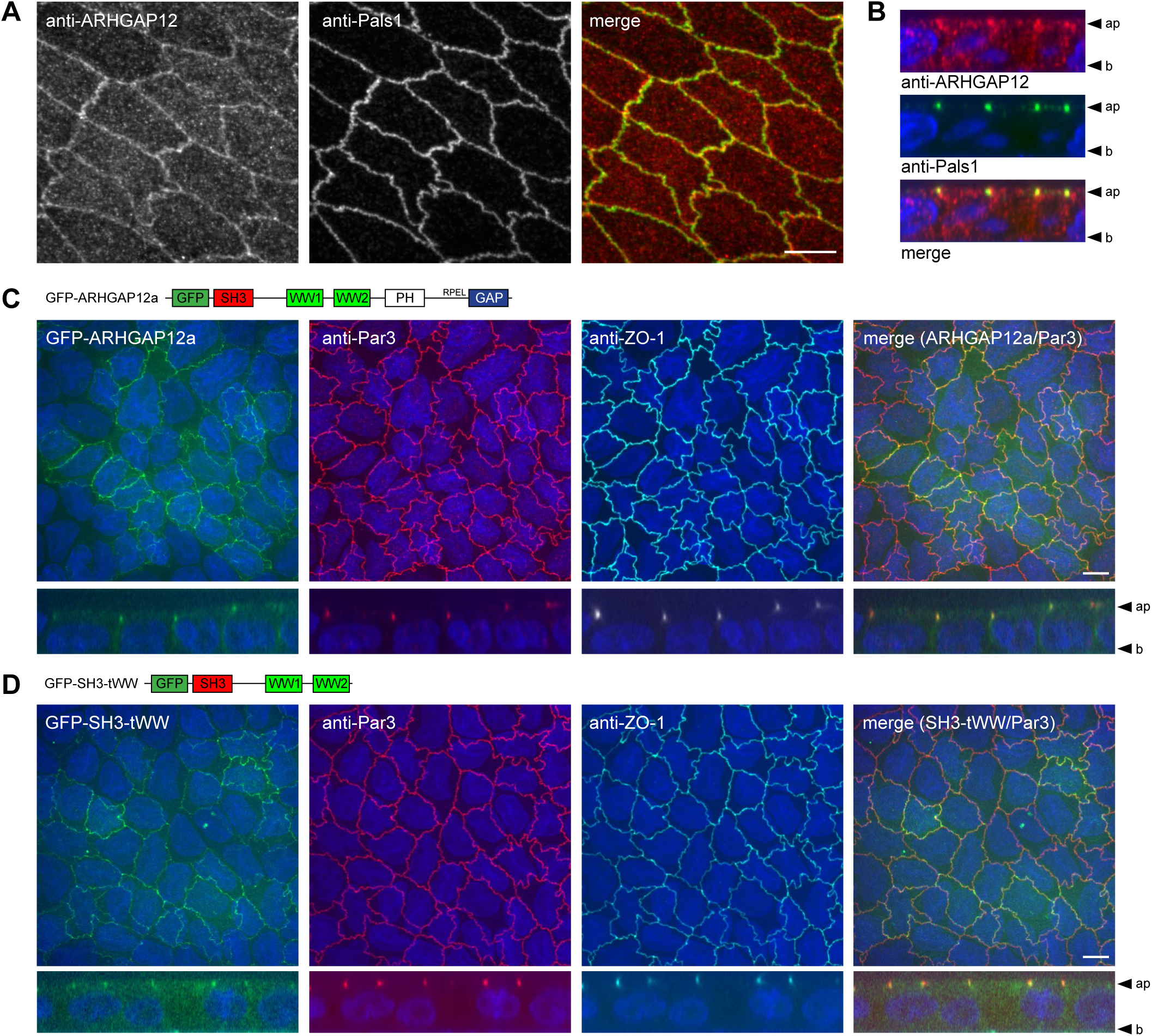
ARHGAP12 localises to tight junctions. (A) Immunofluorescence of MDCK-II cells grown on Transwell filters for 10 days stained with anti-Pals1 and anti-ARHGAP12 antibodies. Apical confocal optical sections are shown. (B) Orthogonal slices (x/z-projections) of the data shown in (A). (C and D) Immunofluorescence of MDCK-II cells grown on Transwell filters for 10 days stably expressing GFP-ARHGAP12a (C) or GFP-SH3-tWW (D). Cells were co-stained with anti-ZO-1 and anti-Par3 antibodies. Maximum intensity projections of confocal z-stacks and orthogonal slices are shown. Scale bars in A-D are 10 µm.

Previous studies demonstrated that the ARHGAP12 SH3 domain binds to a PSTPIPP motif at the C-terminus of ZO-2 (Muller et al., 2020; Okada et al., 2011). To confirm this interaction we immunoprecipitated GFP tagged ZO-2 stably expressed in MDCK-II cells. ARHGAP12 specifically co-immunoprecipitated with ZO-2 but not with ZO-3 (Figure S2A). To test whether the SH3 domain of ARHGAP12 recruits the protein to TJ, MDCK-II cells were transiently transfected with GFP-SH3-tWW, GFP-SH3, GFP-tWW, or GFP alone and analysed by confocal microscopy. GFP-SH3-tWW and GFP-SH3 displayed clear junctional localisation and colocalised with ZO-1. By contrast, GFP and GFP-tWW were distributed diffusely throughout the cytoplasm and cell nuclei, with no evidence of membrane targeting (Figure S2B). Quantitative line scans and Pearson’s correlation analysis confirmed that the SH3 domain is required and sufficient for TJ localisation (Figure S2C and S2D). To address whether the TJ localisation of ARHGAP12 is dependent on ZO proteins, MDCK-II cells stably transfected with Pals1-APEX2-EGFP (Pals1-A2E; as a surrogate for WT cells) were co-cultured with either ZO-1 or ZO-2 knockout (KO) cells. Western blotting (WB) confirmed the complete loss of ZO-1 and ZO-2 in the respective KO cell lines (Figure S2E). ARHGAP12 protein expression was not affected in ZO-1 or ZO-2 KO cells, but junctional ARHGAP12 levels were markedly reduced in both ZO-1 and ZO-2 KO cells (Figure S2F and S2G). Collectively, these data show that ZO proteins mediate the recruitment of ARHGAP12 to TJ, likely via a direct interaction between ZO-2 and the ARHGAP12 SH3 domain.

### ARHGAP12 interacts with N-WASP and the WAVE2 complex

To identify proteins associated with ARHGAP12, GFP-ARHGAP12 transiently transfected into 293T cells was immunoprecipitated using anti-GFP antibodies. Protein bands that were specifically recovered by ARHGAP12 were analysed by Mass Spectrometry. This analysis identified N-WASP and WAVE2 as potential ARHGAP12 binding partners. In addition to the WAVE2 protein, other components of the WAVE regulatory complex (WRC) were identified, including Abi1, Abi2, NAP1 (NCKAP1), CYFIP1 (Sra1) and CYFIP2 (PIR121) (data not shown).

N-WASP and the WRC activate Arp2/3 complex to drive branched actin polymerisation at the cell cortex (Kurisu and Takenawa, 2009). Both N-WASP and the WRC as well as several regulators and effectors of N-WASP and WAVE2 are localised to apical cell junctions (Tan et al., 2020) (Figure S1). To confirm that ARHGAP12 interacts with N-WASP and the WRC, 293T cells were transiently transfected with GFP-ARHGAP12a, GFP-ARHGAP12b, GFP-SH3-tWW, or GFP as a control (Figure 2A). Immunoprecipitations (IPs) using anti-GFP antibodies showed that GFP-ARHGAP12a, GFP-ARHGAP12b and GFP-SH3-tWW indeed interacted with N-WASP, WAVE2 and Abi1 (Figure 2B). An interaction between full-length ARHGAP12 and actin was also identified. This is in line with previous work showing that ARHGAP12 interacts with G-actin through an RPEL motif located just upstream of the GAP domain (Diring et al., 2019). Importantly, Arp2 (a component of the Arp2/3 complex), Mena (a protein involved in actin nucleation and polymerization), and mDia1 (a member of the formin family that nucleate linear actin filaments) were not co-immunoprecipitated. This signifies a specific interaction between ARHGAP12 and N-WASP and WAVE2. To ascertain that these interactions take place in MDCK-II cells, we performed IPs from MDCK-II cells stably transfected with GFP-ARHGAP12 variants. As expected, N-WASP and WAVE2 interacted with full-length ARHGAP12 as well as with the SH3-tWW fragment, whilst actin was only recovered with full-length ARHGAP12 (Figure 2C). ZO-2 and ZO-1 were also co-immunoprecipitated, both by full-length ARHGAP12 and by the N-terminal SH3-tWW domain fragment. By contrast, several activators and effectors of N-WASP or WAVE2 (TRIP10/CIP4, IRSp53, TOCA-1, and TIAM1) as well as the TJ proteins Par3 and claudin-1 did not co-IP with ARHGAP12. We concluded that the SH3-tWW module is sufficient to interact with ZO-2 and ZO-1 and the N-WASP and WAVE2 complexes in MDCK-II cells.

**Figure 2:**
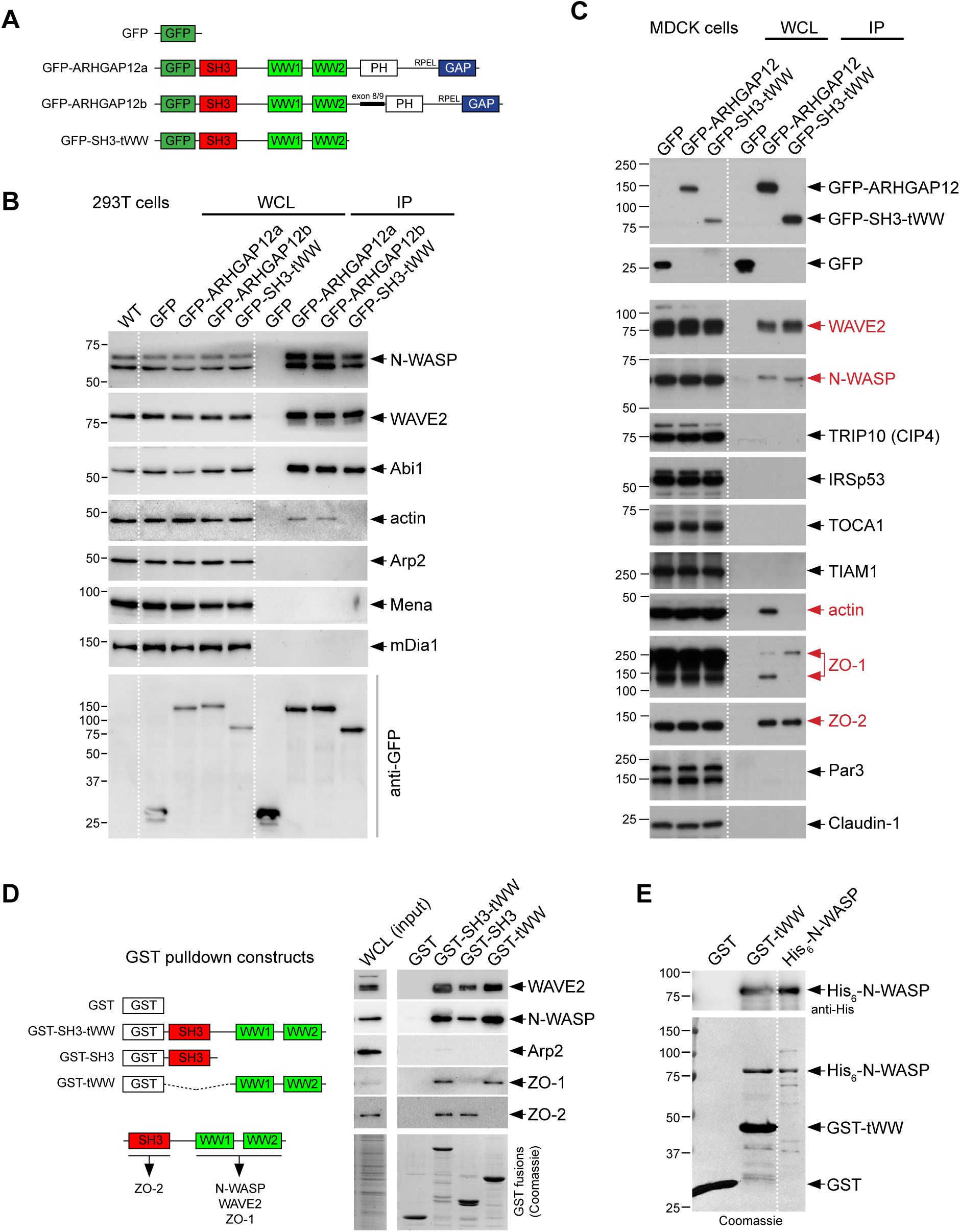
ARHGAP12 interacts with N-WASP and the WAVE2 complex. (A) Domain organisation of ARHGAP12 constructs used in B and C. (B) WB analysis of anti-GFP IPs from 293T cell lysates. Cells were transiently transfected with GFP-ARHGAP12a, GFP-ARHGAP12b, GFP-SH3-tWW or GFP as a control. (C) WB analysis of anti-GFP IPs from MDCK-II cell lysates. Lysates were derived from MDCK-II cell lines stably expressing GFP-ARHGAP12a, GFP-SH3-tWW, or GFP as a control (D) WB analysis of GST pulldowns. Purified GST-ARHGAP12 protein fragments were incubated with MDCK-II cell lysate. Coomassie staining shows the loading control for the GST pulldown. (E) GST pulldown of purified GST-tWW with purified full-length His_6_-N-WASP. A Coomassie-stained gel and a WB is shown.

To address the specific contributions of the SH3 and tWW domains we performed GST pulldowns (Figure 2D). Purified GST fusion proteins were coupled to glutathione sepharose and incubated with MDCK-II cell lysates. Both the SH3 and the tWW domain precipitated WAVE2 and N-WASP from cell lysates, with the tWW domain being a more effective binder than the SH3 domain. Interestingly, whilst the SH3 domain specifically pulled down ZO-2, as expected, ZO-1 was specifically recovered by the tWW domain. To test whether the interaction between the tWW domain and N-WASP and WAVE2 is direct, GST-tWW was incubated with purified full-length proteins of N-WASP (His_6_-N-WASP) or WAVE2 (His_6_-WAVE2). Whilst N-WASP efficiently interacted with GST-tWW (Figure 2E), the purified WAVE2 protein did not (data not shown). This indicates a direct and specific interaction between the tWW domain of ARHGAP12 and N-WASP. On the contrary, the interaction with the WRC observed in our IPs does not appear to be mediated by the WAVE2 protein. Because N-WASP interacted directly with ARHGAP12 and regulates actin assembly at TJ as well as TJ permeability (Garber et al., 2018; Kalailingam et al., 2017; Kovacs et al., 2011; Van Itallie et al., 2015), we focused on exploring the functional significance of this interaction.

### The ARHGAP12 tWW domain interacts with the N-WASP PRD in a multivalent manner

Based on the above we concluded that the SH3 domain recruits ARHGAP12 to TJ, whilst the tWW domain mediates an interaction with N-WASP. WW domains bind to at least four different classes of poly-proline motifs: PPxY, PPLP, PPR and pSP motifs (Macias et al., 2002). We noticed that the N-WASP PRD contains five highly conserved PPPPPxR motifs (from now on referred to as PPxR in short) (Figure 3A and Figure S3A and S3B). Structural predictions using AlphaFold suggested that each of these PPxR motifs are recognized by the ARHGAP12 tWW domain (Figure 3B and Figure S3C). Interestingly, it was invariably the WW2 domain that was predicted to engage with the poly-proline stretch preceding the conserved arginine. In line with published WW domain structures, aromatic amino acids in the WW2 domain, in particular Y129, Y131, and W140 (numbering is based on the tWW domain construct), were predicted to form a hydrophobic groove that accommodated the poly-proline helix.

**Figure 3:**
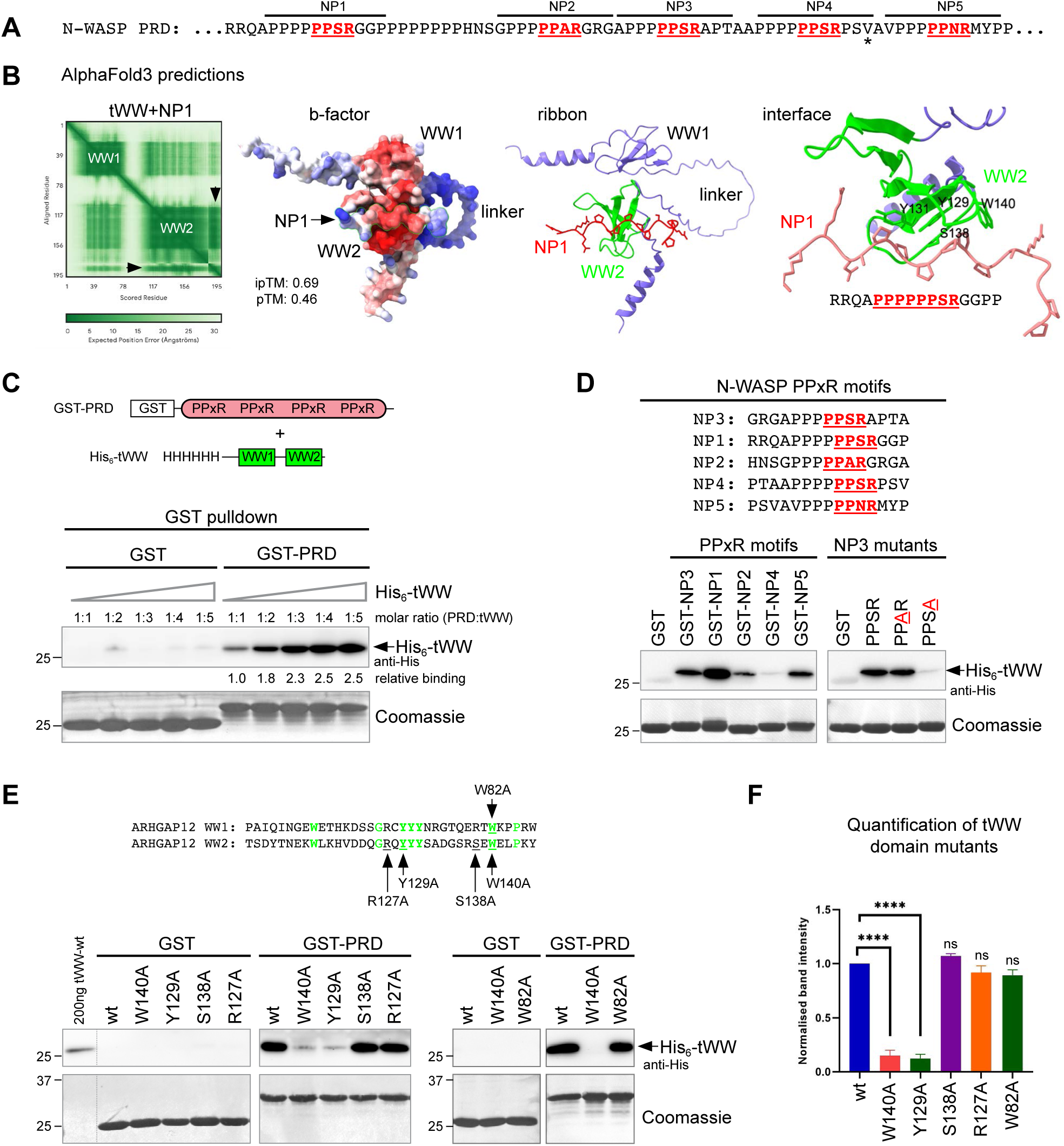
The ARHGAP12 tWW domain interacts directly with PPxR repeats in the N-WASP PRD. (A) Amino acid sequence of the human N-WASP PRD indicating the five conserved PPxR repeats (NP1-NP5). * indicates the C-terminus of the PRD construct used in C and E. (B) AlphaFold3 prediction of the tWW domain of ARHGAP12 bound to the NP1 peptide. From left to right: 1) Predicted Aligned Error (PAE) plot showing that AlphaFold accurately predicts the WW1 and WW2 domain folds. The NP1 peptide is indicated by arrowheads. The ipTM (0.69) and pTM (0.46) values signify that the overall prediction of the structure is of medium to high confidence. 2) Surface representation of the predicted complex shown in b-factor colour code. Red: high confidence, blue: low confidence. 3) Ribbon representation of the complex. 4) Binding interface of the WW2 domain and the NP1 peptide. Key aromatic amino acids (Y129, Y131, W140) of the hydrophobic groove in the WW2 domain are shown. (C) GST pulldowns of the GST-PRD construct with purified His_6_-tWW. Increasing amounts of His_6_-tWW were incubated with GST-PRD or GST as a control. The membrane was stained with Coomassie to serve as a loading control. The relative binding efficiencies were determined by densitometry. (D) Left: GST pulldowns of GST-PPxR motif fusion proteins with purified His_6_-tWW. Right: GST pulldowns of the wild-type GST-NP3 fusion protein and the GST-NP3 S/A or R/A mutants with purified His_6_-tWW. The membrane was stained with Coomassie to serve as a loading control. (E) GST pulldowns of GST-PRD with the wild-type tWW domain protein or tWW domain mutants with point mutations in the WW2 or WW1 domains as indicated. (F) Quantification of the data shown in (E). **** = p<0.0001 (n=3; One-Way ANOVA).

To test the validity of these predictions we first asked whether the tWW domain indeed interacts with the PRD of N-WASP. To this end, we generated a GST fusion protein of the N-terminal ∼60 amino acids of the N-WASP PRD. This construct (GST-PRD) and the His-tagged tWW domain (His_6_-tWW) were expressed in *E.coli* and purified using affinity chromatography and Size Exclusion Chromatography (Figure S4A). Mass photometry (Figure S4B) and Dynamic Light Scattering (not shown) showed that the purified His_6_-tWW protein predominantly exists as a monomer in solution, whilst the GST-PRD protein formed a dimer. GST pulldowns showed that the tWW domain interacts directly with the N-WASP PRD in a concentration-dependent manner (Figure 3C). This demonstrates that the N-terminal ∼60 aa of the N-WASP PRD are sufficient for the interaction with the ARHGAP12 tWW domain.

Next we analysed whether the tWW domain recognizes the conserved PPxR repeats in the N-WASP PRD. To this end, we designed and purified a battery of recombinant proteins in which 12-15 aa short proline-rich peptides derived from the N-WASP PRD were fused to GST (Figure 3D and Figure S4C). Indeed, all five PPxR peptides of the N-WASP PRD interacted with the tWW domain in GST pulldowns (Figure 3D). The NP1 peptide appeared to be the most efficient binder, whilst the NP4 peptide was the least effective. Importantly, mutation of the arginine in the PPxR motif of NP3 (R/A mutant) greatly diminished the binding to the tWW domain (Figure 3D). In addition, several N-WASP PRD peptides of similar length and proline content but lacking a PPxR motif did not interact with the tWW domain (Figure S4C). We concluded that the ARHGAP12 tWW domain interacts specifically with all five PPxR repeats in the N-WASP PRD.

To test whether the tWW/PRD interaction is mediated specifically by the WW2 domain, we introduced point mutations in the hydrophobic grooves of the WW1 and WW2 domains. Mutation of Y129 or W140 in the WW2 domain to alanine (Y129A and W140A) completely abolished the binding to the N-WASP PRD, whereas mutations of R127 or S138 had no effect (Figure 3E and 3F). Interestingly, a W82A mutation in the WW1 domain did not affect the interaction with the tWW domain. This demonstrates that the WW2 domain mediates the interaction with the N-WASP PRD.

The data described above suggested that the five PPxR motifs in the N-WASP PRD constitute a multivalent docking site for the tWW domain. To test this, and to measure the binding affinities between the tWW domain and the PPxR repeats, we performed Isothermal Titration Calorimetry (ITC) (Figure 4 and Figure S4D). N-WASP peptides containing one, two, or three PPxR repeats were injected into the tWW domain. Figure 4B shows representative ITC heat signatures and binding curves for each peptide, whilst Figure 4C shows the average affinities (K_d_), stoichiometries (N sites), and entropies (-Tι1S) based on two independent measurements. The N-WASP peptide NP1 (one PPxR repeat) interacted with the tWW domain with a K_d_ of ∼23 µM and a peptide/tWW domain stoichiometry of ∼1:1 (Figure 4C). Importantly, an NP1 peptide in which the arginine was mutated to alanine (R/A mutant) completely abolished the binding to the tWW domain (Figure S4D. In addition, a peptide derived from Mena containing a PPLP motif did not bind to the tWW domain in ITC (Figure S4D). Extending the NP1 peptide to NP1/2 (two PPxR repeats) increased the affinity of the interaction (K_d_ ∼14 µM) and resulted in a ∼1:2 molar interaction, i.e. two tWW domains were bound to one peptide (Figure 4C). Next we tested two long overlapping N-WASP peptides, each containing three PPxR repeats. One of which (NP1/2/3) spanned the N-terminal part of the PRD, the other (NP3/4/5) the C-terminal part. Both peptides interacted with the tWW domain in a molar ratio of ∼1:3, i.e. three tWW domains were bound to one peptide. Interestingly, the two peptides differed notably in their affinities to the tWW. Whilst the NP1/2/3 peptide bound to the tWW domain with a K_d_ comparable to that of the NP1/2 peptide (K_d_ of ∼14 µM), the affinity of the NP3/4/5 peptide was considerably lower (K_d_ ∼35 µM) (Figure 4C). This indicates that the N-terminal PPxR repeats provide a higher affinity binding site then the C-terminal ones. Moreover, we observed that the entropy (-TΛS) of the system decreased as the number of binding sites on the peptide increased (Figure 4C). This indicates that presenting multiple PPxR repeats promotes order in the tWW/peptide complex, supporting a multivalent interaction. We concluded that the tWW domain of ARHGAP12 interacts with the N-WASP PRD in a specific and multivalent manner.

**Figure 4:**
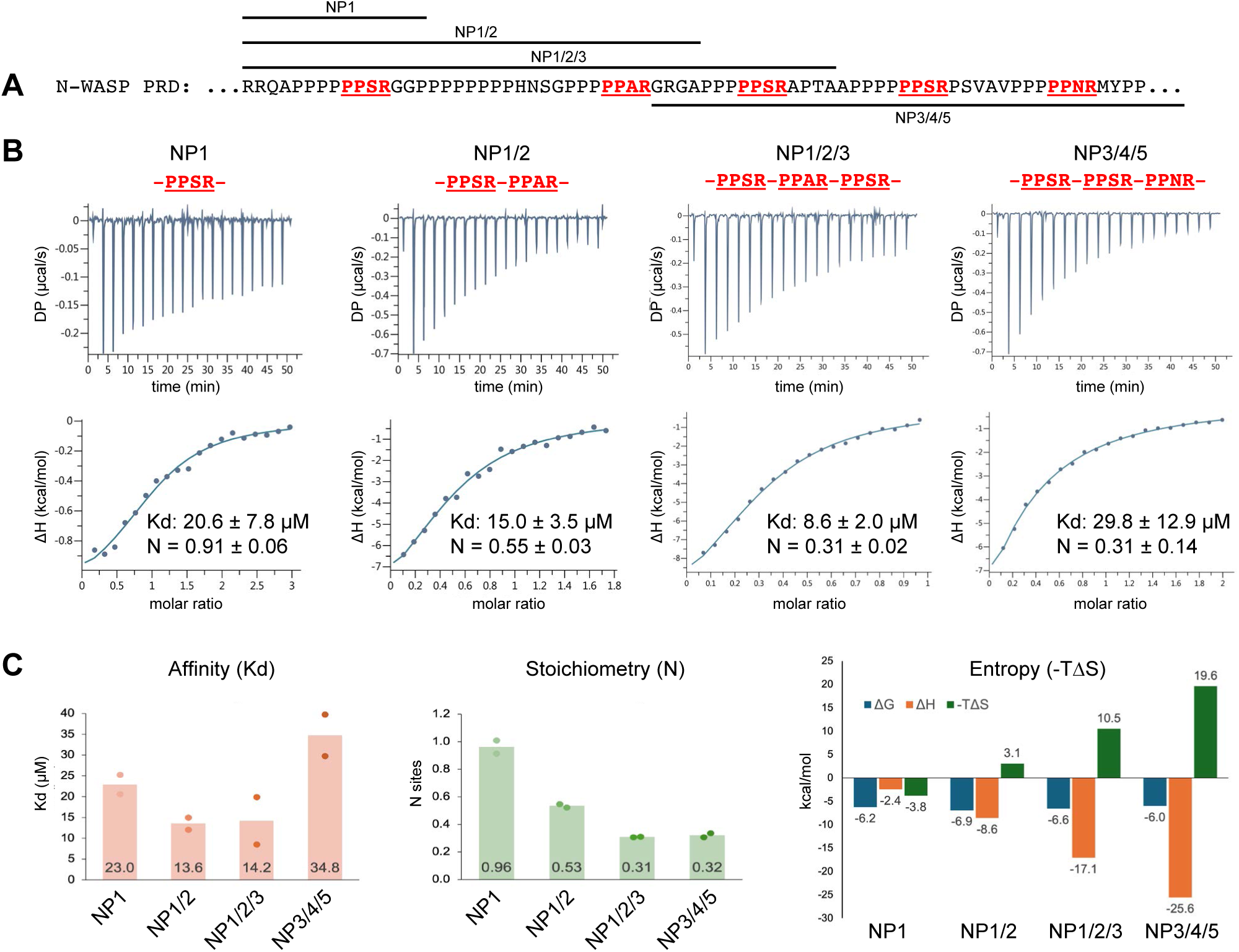
The N-WASP PRD constitutes a multivalent docking site for the ARHGAP12 tWW domain. (A) Amino acid sequence of the human N-WASP PRD indicating the PPxR repeat peptides used in (B). (B) The interaction between the ARHGAP12 tWW domain and the N-WASP PPxR peptides were measured by ITC. The N-WASP peptides NP1, NP1/2, NP1/2/3, or NP3/4/5 (500 µM) were injected into the tWW domain (His_6_-tWW; 50 µM). Representative ITC heat signatures, binding curves, and the corresponding K_d_ (affinity) and N (stoichiometry) values of a representative experiment are shown. (C) ITC data showing the average affinity (K_d_), stoichiometry (N sites), and entropy (-TΔS) of the interaction between the tWW domain and the indicated N-WASP peptides based on two independent measurements.

### The ARHGAP12 tWW domain is sufficient to suppress N-WASP activity in vitro

To address the function of the ARHGAP12/N-WASP interaction we used a TIRF-based in vitro actin polymerisation assay (Figure 5 and Figure S5). This assay monitors branched F-actin assembly in a minimal system composed of fluorescently labelled G-actin, purified Arp2/3 complex, and purified N-WASP protein. Wild-type N-WASP is autoinhibited and therefore does not activate Arp2/3 complex (Figure 5A). However, when the assay is performed with a constitutively active mutant of N-WASP (L235P) that relieves N-WASP autoinhibition, strong F-actin assembly is observed (Figure S5A). Addition of the purified tWW domain did not significantly affect the number or the size of F-actin clusters generated by the N-WASP L235P mutant (Figure S5A). However, we noted that the tWW domain significantly reduced the fluorescence intensity in the centre of the F-actin clusters in a concentration-dependent manner (Figure S5B). This suggests that under these experimental conditions the tWW domain interferes with the formation or stability of N-WASP actin nucleation centres.

**Figure 5:**
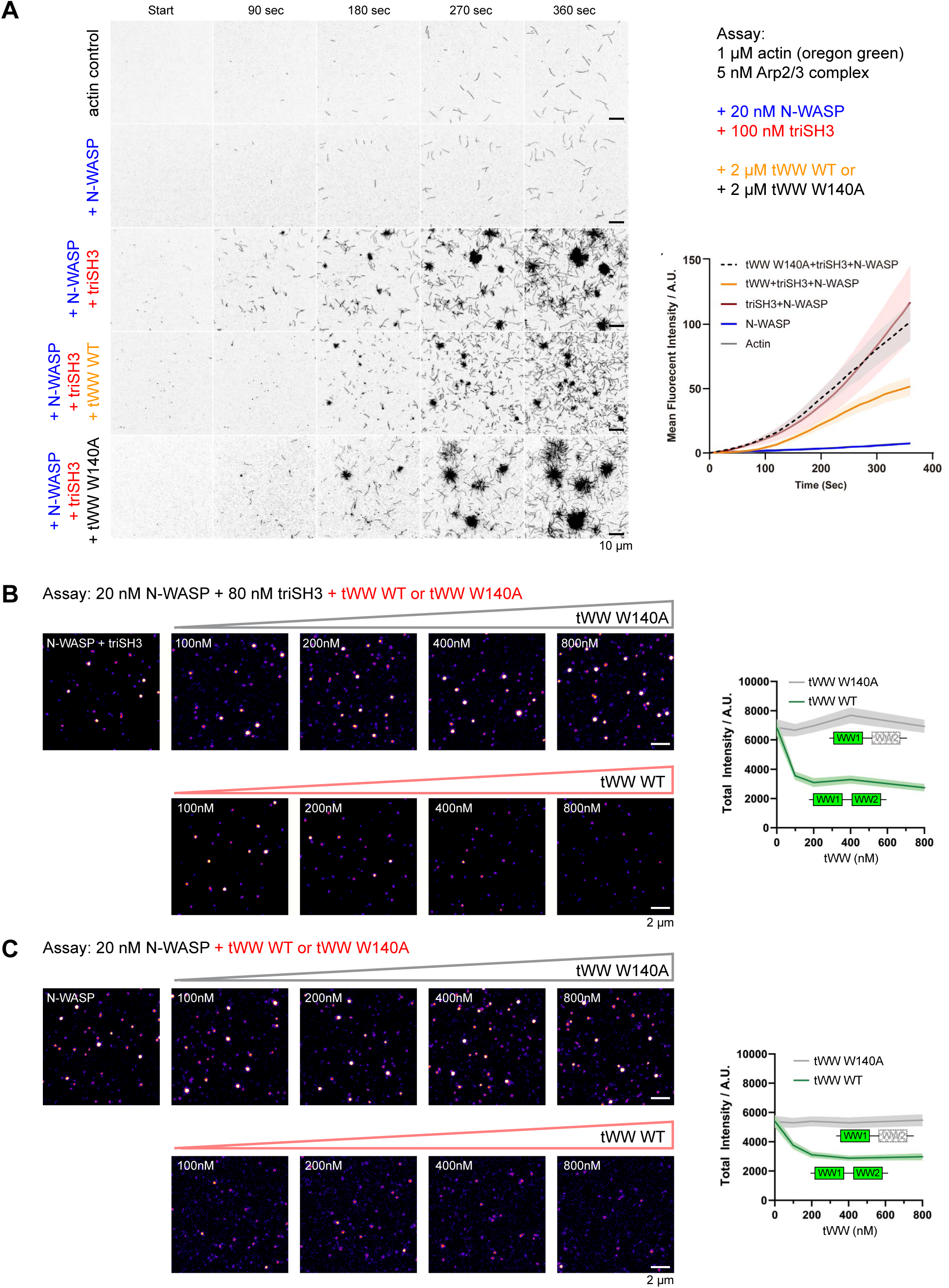
The ARHGAP12 tWW domain suppresses N-WASP-driven Arp2/3 mediated actin assembly and N-WASP oligomerisation in vitro. (A) In vitro actin polymerisation assay. Fluorescently labelled (Oregon green) G-actin, Arp2/3 complex, wild-type N-WASP, and a triSH3 fragment derived from FBP17 were incubated in the absence or presence of the wild-type ARHGAP12 tWW domain or the tWW domain W140A mutant. Actin polymerisation was monitored live using TIRF microscopy. (B and C) Single molecule TIRF imaging. Fluorescently labelled wild-type N-WASP was incubated with (B) or without (C) the triSH3 fragment and increasing concentrations of the wild-type tWW domain or the tWW domain W140A mutant.

The activity of wild-type N-WASP can also be greatly enhanced by oligomeric SH3 domains. Such ligands bind to the N-WASP PRD in a multivalent manner, resulting in N-WASP oligomerisation and increased Arp2/3-mediated actin assembly (Padrick et al., 2008; Padrick and Rosen, 2010). To address whether the tWW domain affects N-WASP activity in the presence of such strong activating signals, we generated an artificial trimeric SH3 protein derived from the SH3 domain of FBP17 (referred to as triSH3). The triSH3 protein self-assembles in solution from SH3 monomers via an N-terminal oligomerisation tag, which mediates the formation of a stable trimeric coiled-coil (Khairil Anuar et al., 2019). When the triSH3 protein was added to wild-type N-WASP, Arp2/3 complex and G-actin, F-actin polymerisation was strongly enhanced compared to control experiments lacking the triSH3 activator (Figure 5A). Interestingly, addition of the purified wild-type tWW domain, but not the W140A tWW mutant, significantly attenuated F-actin assembly in the presence of triSH3 (Figure 5A). This demonstrates that the tWW domain is sufficient to suppress SH3 domain-mediated N-WASP activation and Arp2/3-driven actin assembly in vitro.

Next we asked whether the tWW domain affected triSH3-mediated N-WASP oligomerisation using a TIRF-based single molecule fluorescence microscopy assay. Purified and fluorescently labelled wild-type N-WASP was incubated with the triSH3 protein and increasing concentrations of the wild-type tWW domain or the W140A tWW domain mutant. Intriguingly, the wild-type tWW domain but not the W140A tWW mutant effectively prevented N-WASP oligomerisation (Figure 5B). Full suppression of oligomerisation was already evident at a triSH3/tWW molar ratio of ∼1:1. N-WASP oligomerisation was similarly inhibited in the absence of the triSH3 protein (Figure 5C). This indicates that the ARHGAP12 tWW domain can effectively suppress N-WASP oligomerisation in a GAP-independent manner, even in the presence of trimeric SH3 domain ligands.

### SH3 domains do not effectively compete with the ARHGAP12 tWW domain for PRD binding

We wanted to understand how the tWW domain interferes with SH3 domain-mediated N-WASP activation. SH3 and WW domains recognise similar poly-proline motifs, suggesting signaling crosstalk and competition (Li, 2005; Macias et al., 2002). We therefore analysed how the triSH3 protein and other SH3 domains interact with the N-WASP PRD, and whether they can compete with the tWW domain for PRD binding. The purified triSH3 protein, but not the monomeric FBP17 SH3 domain, efficiently interacted with the N-WASP PRD in pulldown assays (Figure 6A and 6B). Although the triSH3 protein interacted most efficiently with the N-WASP NP3 peptide, all five N-WASP PPxR repeats and a poly-proline peptide lacking a PPxR motif (NP6) were recognised by the triSH3 fragment (Figure 6C). By contrast, both the ARHGAP12 SH3 domain and the TUBA SH3 domain, which is known to interact with the N-WASP PRD (Salazar et al., 2003), specifically interacted with the NP3 peptide (Figure 6D and 6E). This suggests that the triSH3 protein binds promiscuously to poly-proline stretches, whilst the ARHGAP12 and TUBA SH3 domains are far more selective poly-proline binders. To test whether SH3 domains and the tWW domain compete for PRD binding, immobilised GST-PRD was saturated with the tWW domain and subsequently incubated with increasing amounts of the triSH3 fragment. The data showed that the triSH3 domain did not displace the tWW domain from the PRD (Figure 6F). Similarly, neither the ARHGAP12 nor the TUBA SH3 domains interfered with the tWW/PRD interaction in such competition pulldown assays, even when they were present in several fold molar access (Figure 6G). This indicates that the tWW domain of ARHGAP12 utilizes a binding mechanism distinct from that employed by SH3 domain containing PRD ligands. Altogether our data suggest that the tWW domain interferes with N-WASP oligomerisation by occupying SH3 domain binding sites required for N-WASP crosslinking (Figure 6H).

**Figure 6:**
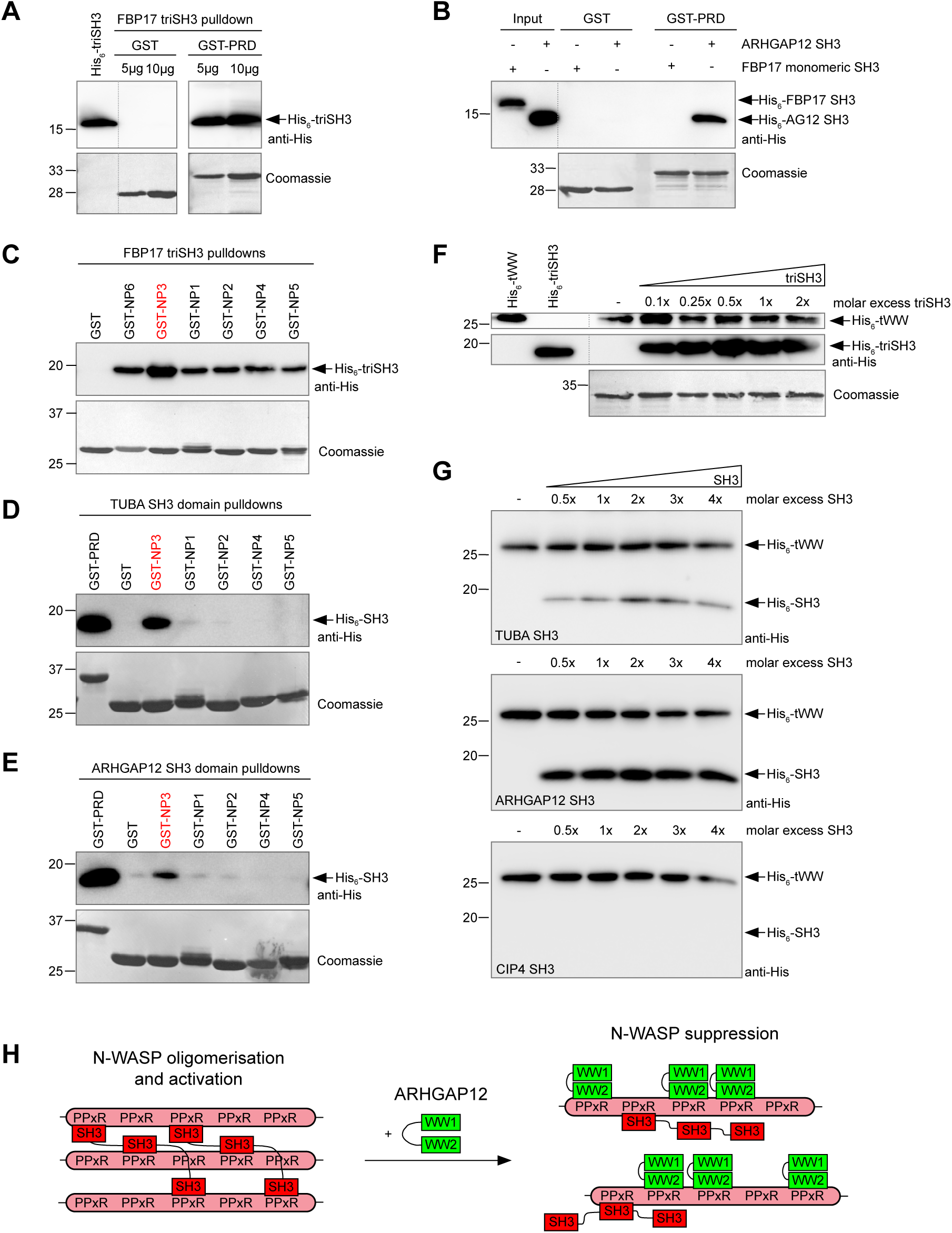
SH3 domains do not effectively compete with the ARHGAP12 tWW domain for binding to the N-WASP PRD. (A) Purified GST or GST-PRD was incubated with purified His_6_-triSH3. (B) Purified GST or GST-PRD was incubated with either the purified FBP17 SH3 domain or the purified ARHGAP12 SH3 domain. (C-E) GST pulldowns of purified GST N-WASP peptide fusion proteins with the FBP17-derived triSH3 fragment (C), the TUBA SH3 domain (D), or the ARHGAP12 SH3 domain (E). GST-NP6 represents a non-PPxR peptide (PPPPPALPSSAP). The sequences of all other peptides are shown in Figure 3D. (F) Competition assay. The GST-PRD was saturated with the purified His_6_-tWW domain, followed by the addition of increasing amounts of the triSH3 fragment. (G) Competition assay. The GST-PRD was saturated with the purified His_6_-tWW domain, followed by the addition of increasing amounts of the TUBA, ARHGAP12, or CIP4 (TRIP10) SH3 domains. Note that the CIP4 SH3 domain does not interact with the N-WASP PRD. (H) Model of how the ARHGAP12 tWW domain interferes with SH3 domain-mediated N-WASP oligomerisation.

### ARHGAP12 colocalises with N-WASP and protrusive F-actin networks in cells

To address the significance of the ARHGAP12/N-WASP interaction in a cellular context, we first analysed the colocalization between ARHGAP12, N-WASP, and F-actin in transfected COS7 and 293T cells (Figure 7 and Figure S6). As expected, both full-length GFP-ARHGAP12 and GFP-SH3-tWW colocalised with F-actin patches and N-WASP in fixed cells (Figure S6). GFP-SH3-tWW and endogenous N-WASP colocalised in discrete puncta at the cell periphery (Figure 7A) and this colocalization was significantly reduced in cells expressing the GFP-SH3-tWW W140A mutant (Figure 7B and 7C). In addition, N-WASP fluorescence intensity was increased ∼2-fold in cells expressing GFP-ARHGAP12 or GFP-SH3-tWW, but was unchanged in cells expressing the GFP-SH3-tWW W140A mutant (Figure 7D). In live cells, GFP-N-WASP and mCherry tagged ARHGAP12 colocalised in dynamic cortical patches that persisted for several minutes (Figure 7E and Movie S1). Interestingly, live cell imaging with the actin probe LifeAct-mCherry showed that the GFP-SH3-tWW fragment localized specifically to peripheral protrusive actin networks and was conspicuously absent from linear actin stress fibres (Figure 7F and 7G and Movie S2). These data indicate that ARHGAP12 interacts with N-WASP in live cells in a WW2 domain-dependent manner.

**Figure 7:**
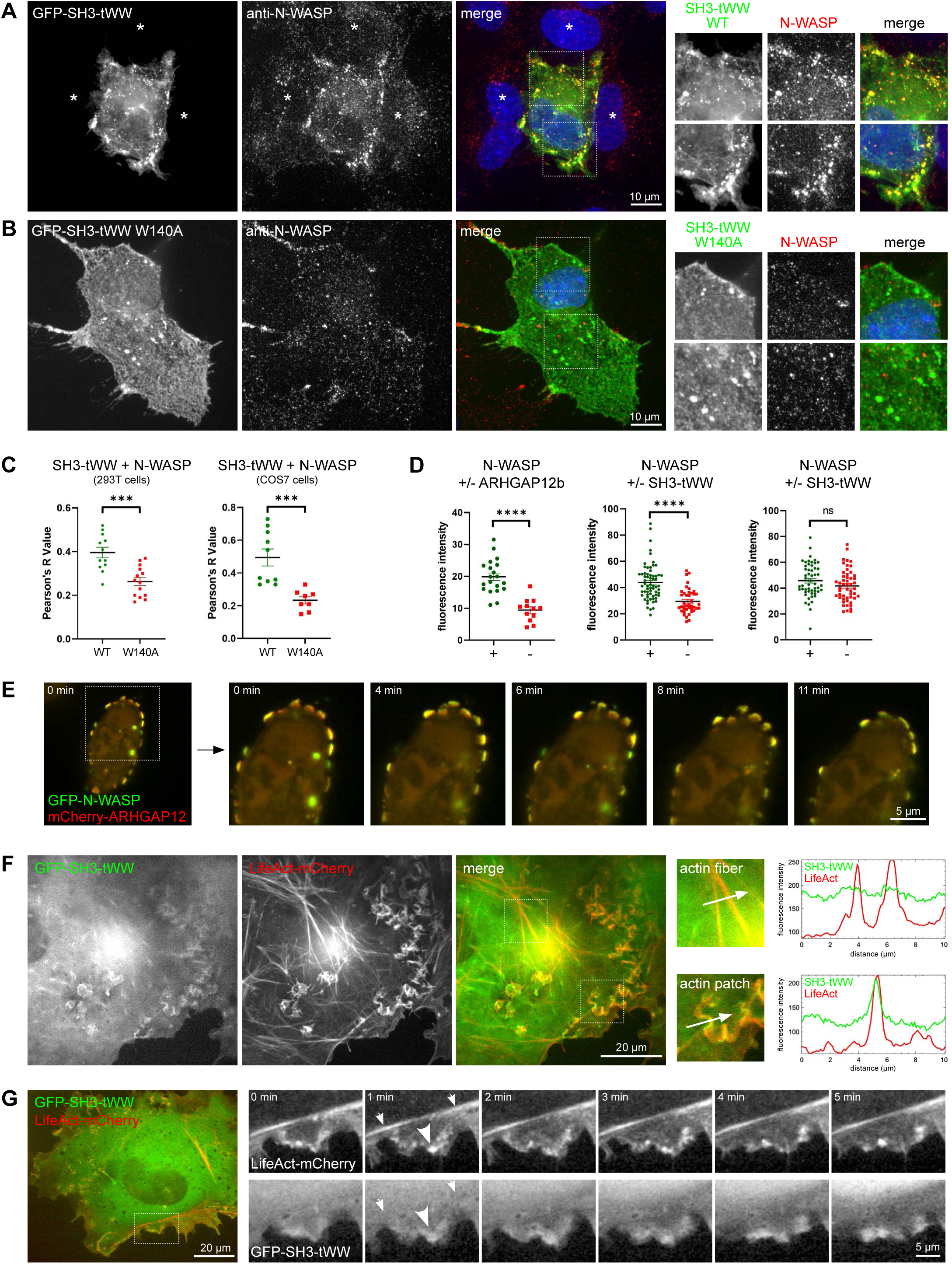
ARHGAP12 colocalises with N-WASP and protrusive F-actin networks. (A and B) COS7 cells transfected with wild-type GFP-SH3-tWW (A) or the GFP-SH3-tWW W140A mutant (B) were fixed, stained with anti-N-WASP antibodies and imaged by confocal microscopy. Note the increase in N-WASP fluorescence intensity in cells transfected with the wild-type SH3-tWW domain compared to untransfected cells (* in A). (C) Colocalisation analysis (Pearson’s correlation) of GFP-SH3-tWW or the GFP-SH3-tWW W140A mutant with endogenous N-WASP in 293T and COS7 cells. *** = p<0.001 (student’s t-test). Error bars indicate the standard error of the mean (SEM). (D) Quantification of N-WASP fluorescence intensity in 293T cells overexpressing GFP-ARHGAP12, GFP-SH3-tWW, or the GFP-SH3-tWW W140A mutant. “+” = transfected cells, “-“ = untransfected cells. **** = p<0.0001 (student’s t-test). Error bars indicate the standard error of the mean (SEM). (E) 293T cells co-transfected with mCherry-ARHGAP12 and GFP-N-WASP were imaged live using spinning disk microscopy at a frame rate of 10 sec. (F) Confocal micrographs of live COS7 cells co-transfected with GFP-SH3-tWW and LifeAct-mCherry. The fluorescence intensity of GFP-SH3-tWW and LifeAct-mCherry in actin fibers and actin patches was measured with line scans (indicated as arrows). (G) COS7 cells co-transfected with GFP-SH3-tWW and LifeAct-mCherry were imaged live using spinning disk microscopy at a frame rate of 10 sec. Note that GFP-SH3-tWW localises to cortical actin networks (large arrowheads) but not to linear actin fibers (small arrowheads).

### ARHGAP12 controls TJ leak pathway permeability and junctional tension via N-WASP

Our data led us to speculate that ARHGAP12 suppresses N-WASP at TJ. To test this, we generated ARHGAP12 MDCK-II KO cells using CRISPR/Cas9. Genomic DNA sequencing revealed distinct mutations in all three ARHGAP12 KO clones selected (clones F9, F21, and F23). All mutations resulted in a premature stop codon not far downstream of the respective guide RNA target sites (Figure S7A). WB and immunofluorescence microscopy confirmed that the expression of ARHGAP12 was abolished in all three KO lines (Figure S7B and S7C). Loss of ARHGAP12 did not affect the expression levels of N-WASP, WAVE2, G-actin, or Arp2. The levels of ZO-1, ZO-2, several claudins and occludin were also unchanged (Figure S7B) and TJ morphology appeared normal when cells were cultured to full confluence (Figure S7C and S7D). We concluded that inactivation of *ARHGAP12* does not noticeably perturb the expression or subcellular localization of TJ proteins.

To test whether ARHGAP12 regulates TJ formation, we performed calcium switch assays. Calcium depletion resulted in the complete loss of cell-cell junctions in both wild-type (WT) and ARHGAP12 KO cells, as judged by ZO-1 staining (Figure S8). In WT cells, TJ were fully re-established 4-6 hours post calcium addition, as expected (Figure 8A and Figure S8B). By contrast, ARHGAP12 KO lines were clearly defective in forming a continuous TJ belt (Figure 8BA and Figure S8C and S8D). Quantification of the average junction length revealed a significant defect in junction reformation in both ARHGAP12 KO clones within the first 6 hours of calcium addition (Figure 8B). Even after 24 hours the cell-cell contacts of KO cells appeared discontinuous (Figure 8A and Figure S8C and S8D). To support a role for ARHGAP12 in the formation of a functional TJ belt, we monitored the establishment of trans-epithelial electrical resistance (TEER). MDCK-II WT and ARHGAP12 KO cells were seeded on Transwell filters and TEER was measured during the first 7 days in culture. Whilst WT cells established peak TEER values at ∼24 hrs, ARHGAP12 KO cells reached the TEER peak at ∼48 hrs (Figure 8C). In addition, in WT cells steady state TEER values were established at ∼48 hrs, whereas ARHGAP12 KO cells reached steady state TEER values only after ∼5-6 days in culture. From this point onwards, no significant difference in TEER was observed between WT and KO cells. We concluded that ARHGAP12 promotes TJ formation and the establishment of a normal pore pathway. However, paracellular ion flux is unaffected once TJ have matured.

**Figure 8:**
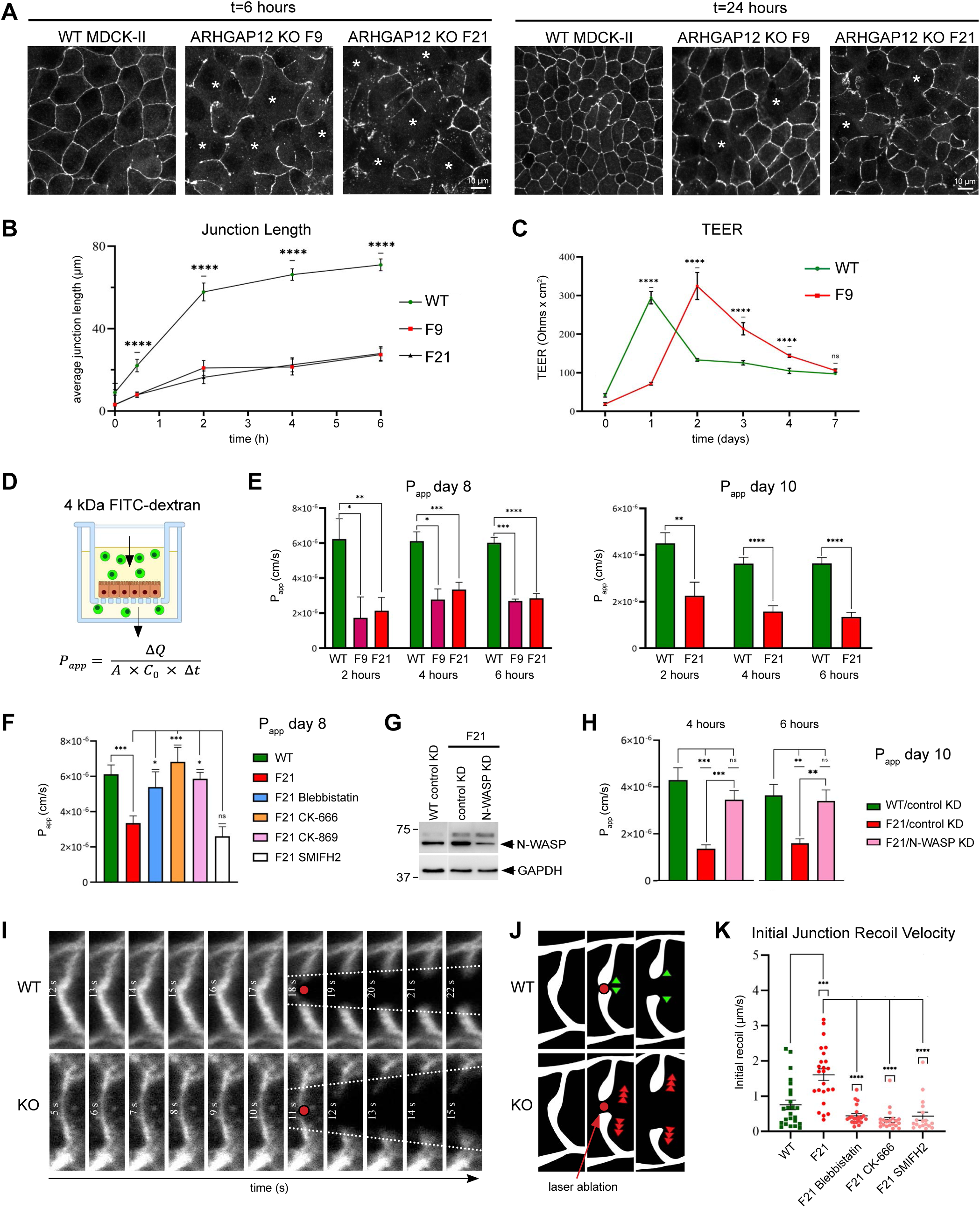
ARHGAP12 promotes TJ formation and controls the TJ leak pathway and junctional tension via the actin cytoskeleton. (A) Calcium switch assay of MDCK-II WT and MDCK-II ARHGAP12 KO cells (clones F9 and F21). Cells were fixed at different time points post calcium addition and processed for immunofluorescence using anti-ZO-1 antibodies. Representative areas at 6 hrs and 24 hrs post calcium addition are shown. Note the discontinuous TJ (*) in ARHGAP12 KO cells. (B) Quantification of the average junction length in MDCK-II WT and ARHGAP12 KO cells (clones F9 and F21) subjected to calcium switch. **** = p<0.0001 (student’s t-test; n=3; with 50-70 cells per replicate). Error bars indicate the standard error of the mean (SEM). (C) TEER measurements of MDCK-II WT and ARHGAP12 KO cells (clone F9) cultured on Transwell filters. TEER was measured from day 0 to day 8 after seeding. *** = p<0.001, **** = p<0.0001 (student’s t-test; n=3). Error bars represent standard deviation (SD). (D) The apparent permeability (P_app_) of MDCK monolayers to 4 kDa FITC dextran grown on Transwell filters was calculated using the formula shown. ΔQ is the amount of dextran detected in the basolateral side (mg), A is the surface area of the insert (cm^2^), C_0_ is the initial concentration in the apical compartment (mg/ml), and Δt is the time of the experiment (s). (E) P_app_ of MDCK-II WT and ARHGAP12 KO cells (clones F9 and F21) at 8 and 10 days on filters. P_app_ was measured at 2, 4, and 6 hrs after addition of 4 kDa FITC-dextran. * = p<0.05; ** = p<0.01; *** = p<0.001; **** = p<0.0001 (student’s t-test; n=3-5). Error bars indicate the standard error of the mean (SEM). (F) P_app_ of MDCK-II WT and ARHGAP12 KO F21 cells with and without drug treatment (50 µM blebbistatin, 200 µM CK-666, 200 µM CK-869, or 10 µM SMIFH2). P_app_ was measured at day 8 on filters and 4 hrs after simultaneous addition of the drug and 4 kDa FITC-dextran. * = p<0.05; *** = p<0.001 (student’s t-test; n=3). Error bars indicate the standard error of the mean (SEM). (G) Representative WB of N-WASP protein levels in wild-type MDCK cells stably transfected with a control shRNA and ARHGAP12 F21 KO MDCK cells stably transfected with a control shRNA or an N-WASP shRNA. (H) P_app_ of control shRNA MDCK-II cells, ARHGAP12 KO F21 control shRNA cells, and ARHGAP12 KO F21 cells transfected with N-WASP shRNA. P_app_ was measured at day 10 on filters, 4 and 6 hrs after addition of 4 kDa FITC-dextran. ** = p<0.01; *** = p<0.001 (student’s t-test; n=3). Error bars indicate the standard error of the mean (SEM). (I) Sequential still images of a representative laser ablation experiment. Cells were grown to confluency and stained with CellMask™ to visualize the plasma membrane. Laser ablation was performed on the most apical aspect of the lateral membrane. (J) Schematic of the laser ablation experiment. (K) Quantification of the initial junction recoil velocity in MDCK-II WT and ARHGAP12 KO F21 cells. Cells were either left untreated or treated with 50 µM blebbistatin, 200 µM CK-666, or 10 µM SMIFH2 for 4 hours. *** = p<0.001; **** = p<0.0001 (student’s t-test). Error bars indicate the standard error of the mean (SEM).

Previous studies have implicated N-WASP and Arp2/3 complex in the control of the TJ leak pathway (Garber et al., 2018; Van Itallie et al., 2015). We therefore explored whether ARHGAP12 regulates the transport of macromolecules across TJ. To this end, MDCK-II WT and ARHGAP12 KO cells were cultured on Transwell filters for 8-10 days. 4 kDa FITC-dextran was administered to the apical compartment and samples were collected from the basal compartment after 2, 4, or 6 hours to measure the apparent permeability (P_app_) of the monolayer (Figure 8D). Interestingly, ARHGAP12 KO cells demonstrated a significant decrease in dextran permeability compared to WT cells at all time points tested (Figure 8E). Next we asked whether this increased tightness of ARHGAP12 KO monolayers was caused by changes in the junction-associated actin cytoskeleton. Treatment of ARHGAP12 KO cells with the myosin-II inhibitor blebbistatin or the Arp2/3 inhibitors CK-666 and CK-869 increased dextran permeability to levels comparable to that observed in WT cells (Figure 8F). By contrast, inhibition of formin activity with SMIFH2 had no effect on dextran permeability (Figure 8F). Finally, to test whether the ARHGAP12 permeability phenotype was caused by increased N-WASP activity, we stably downregulated N-WASP in ARHGAP12 KO cells using a specific shRNA. As a control, a non-targeting shRNA was transfected into ARHGAP12 and WT MDCK cells. The shRNA reduced total N-WASP protein levels by ∼50% (Figure 8G). Interestingly, knockdown of N-WASP in ARHGAP12 KO cells significantly elevated 4 kDa dextran permeability, reaching permeability values comparable to those of WT control cells (Figure 8H). We concluded that ARHGAP12 regulates the TJ leak pathway via N-WASP and Arp2/3-mediated junctional actin assembly.

N-WASP and Arp2/3 complex have been shown to regulate junctional tension, and changes in junctional tension have been implicated in the control of the leak pathway (Monaco et al., 2021; Van Itallie et al., 2015; Wu et al., 2014). To test whether junctional tension is affected in ARHGAP12 KO cells, we performed laser ablation experiments. MDCK-II WT and ARHGAP12 KO cells were grown to confluency and stained live with CellMask to outline the plasma membrane. Laser ablations were carried out at apical cell junctions using a 355 nm laser (Figure 8I and 8J and Movie S3). The initial displacement (recoil velocity) of tricellular contacts was calculated as described in the Materials and Methods. We observed a substantial increase in the initial junctional recoil velocity in ARHGAP12 KO cells (1.606 ± 0.165 µm/s) when compared to WT cells (0.754 ± 0.136 µm/s) (Figure 8K). Inhibition of myosin-II (blebbistatin), Arp2/3 (CK-666), or formin (SMIFH2) activities in ARHGAP12 KO cells reduced recoil velocity to 0.450 ± 0.059 µm/s, 0.332 ± 0.067 µm/s, and 0.440 ± 0.114 µm/s, respectively. This demonstrates that the loss of ARHGAP12 considerably increases junctional tension via the deregulation of the actin cytoskeleton. Taken together our data indicate that ARHGAP12 dampens F-actin assembly at TJ to suppress junctional tension, which in turn promotes TJ leak pathway permeability.

## Discussion

Previous work has demonstrated that ARHGAP12 limits Rac or Cdc42 activities to suppress F-actin assembly (Ba et al., 2016; Diring et al., 2019; Gentile et al., 2008; Schlam et al., 2015). However, the functions of the N-terminal SH3 and tWW domains of ARHGAP12 had not been addressed. In addition, although ARHGAP12 has been shown to localise to the apical junctional complex in various epithelial tissues, including the pancreas, the small intestine, the liver, and the kidney (Matsuda et al., 2008), its functions in epithelial cells had not been explored. Here we demonstrate that ARHGAP12 is recruited to TJ via ZO proteins to regulate the epithelial paracellular permeability barrier. The data indicate that ARHGAP12 suppresses N-WASP-mediated branched actin assembly at TJ to dampen junctional tension, which in turn promotes macromolecular transport via the TJ leak pathway (Figure 9).

**Figure 9:**
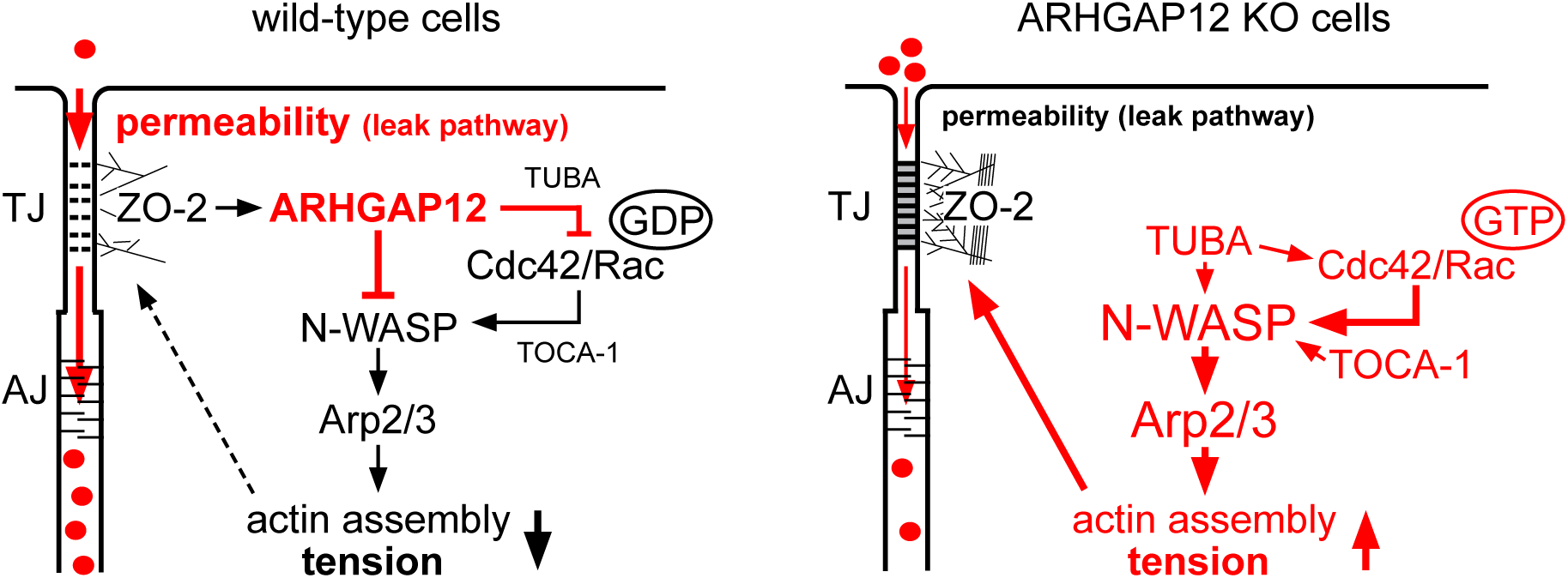
Model of how ARHGAP12 controls the TJ leak pathway. ARHGAP12 is recruited to TJ via ZO-2 and suppresses N-WASP via a direct inhibitory interaction with the PRD and by inhibiting Rac/Cdc42-GTP loading. This dampens F-actin assembly via Arp2/3 complex, decreases junctional tension, and promotes leak pathway permeability for small macromolecules in the 4 kDa size range. In ARHGAP12 KO cells, N-WASP activity is enhanced, resulting in increased F-actin polymerisation and tension, and a tighter barrier for small macromolecules. We speculate that ARHGAP12 interferes with the function of SH3 domain containing N-WASP effectors and activators, such as TOCA-1 and TUBA.

### The ARHGAP12 tandem WW domain interacts with and suppresses N-WASP

We demonstrate that the ARHGAP12 tWW domain interacts specifically and directly with several highly conserved PPxR motifs in the PRD of N-WASP. tWW domains are versatile and interact with proline-rich ligands via diverse mechanisms and with affinities ranging from a few nanomolar to hundreds of micromolar (Dodson et al., 2015; Lin et al., 2019; Webb et al., 2011; Wenz et al., 2022). We show that single point mutations (Y129 and W140) in the hydrophobic groove of the WW2 domain completely abolish the binding to the N-WASP PRD, whereas an equivalent mutation in the WW1 domain (W82A) has no effect. This indicates that a functional WW2 domain is essential for PRD binding. Although the function of the WW1 domain is currently unclear, we speculate that it might act as a chaperone for the WW2 domain or increase the binding affinity or specificity of the tWW domain module, as has been demonstrated for other tWW domains (Dodson et al., 2015). Indeed, the ARHGAP12 tWW domain exhibits a remarkable preference for PPxR repeats. Poly-proline peptides lacking a PPxR motif did not interact with the tWW domain, neither in GST pulldowns nor in ITC. In addition, the actin binding proteins Mena and mDia1, which contain PRDs with several PPLP repeats, did not co-immunoprecipitate with ARHGAP12. Altogether the data suggest that the ARHGAP12 WW2 domain utilizes a binding mechanism akin to that of the WW domains of FBP30, FBP21, and FE65, which engage with proline/arginine motifs featuring an optimal consensus sequence of PPPPR (Bedford et al., 2000). In fact, the ARHGAP12 tWW domain exhibited affinities towards PPxR peptides comparable to those reported for the FBP30 WW domain and its ligand (Bedford et al., 2000).

Interestingly, ITC measurements demonstrated that the ARHGAP12 tWW domain interacts with the N-WASP PRD in a multivalent fashion. Presenting several PPxR repeats also increased the affinity of the interaction, suggesting cooperativity between individual binding events. Multivalency and cooperativity play an important role in SH3 domain-mediated N-WASP activation (Case et al., 2019; Padrick et al., 2008; Padrick and Rosen, 2010). Indeed, we provide evidence that the multivalent interaction between the tWW domain and the N-WASP PRD is functionally relevant. Using single molecule imaging and an in vitro actin polymerisation assay we find that the tWW domain is sufficient to block SH3 domain-mediated N-WASP oligomerisation and N-WASP-driven actin assembly via Arp2/3 complex. In addition, full-length ARHGAP12 and the SH3-tWW fragment efficiently colocalised with N-WASP and protrusive F-actin networks in cells. Both the colocalization with N-WASP and the suppression of F-actin assembly in vitro was dependent on a functional WW2 domain. This indicates that the multivalent nature of the tWW/PRD interaction permits ARHGAP12 to suppress N-WASP in a GAP domain-independent manner.

It is well established that multimeric SH3 domains promote N-WASP oligomerisation and phase separation and thereby greatly enhance Arp2/3-driven actin assembly (Banjade et al., 2015; Case et al., 2019). The TOCA family members FBP17, CIP4, and TOCA-1 as well as the Cdc42 GEF TUBA all interact with the N-WASP PRD via their SH3 domains and thereby activate N-WASP (Ho et al., 2004; Kovacs et al., 2006). The underlying binding mechanisms, however, are often not well understood. In agreement with previous work (Salazar et al., 2003), we find that the TUBA SH3 domain interacts specifically with the NP3 PPxR peptide. The ARHGAP12 SH3 domain also interacted preferentially with this region of the N-WASP PRD, whereas the FBP17 triSH3 fragment bound promiscuously to several PPxR and non-PPxR containing peptides. Interestingly, neither monomeric SH3 domains nor the triSH3 protein were able to displace the tWW domain from the N-WASP PRD in pulldown assays. We propose, therefore, that the tWW domain interferes with the oligomerisation of N-WASP by reducing (or “diluting”) the number of SH3 domain binding sites on the PRD. It is also possible that the tWW and SH3 domains bind to overlapping poly-proline motifs but in opposite orientations (Li, 2005; Macias et al., 2002; Wenz et al., 2022). This might interfere with SH3 domain mediated N-WASP oligomerisation through a steric effect or via conformational changes in the poly-proline helix.

Taken together, our data suggest that ARHGAP12 acts as a “sponge” that sequesters N-WASP in an inactive state. This provides a mechanism to prevent N-WASP activation by monomeric or non-specific SH3 domains and ensures that N-WASP is activated only when a certain local threshold of multimeric SH3 domains has been reached. To our knowledge this is the first demonstration that WW domains can suppress SH3 domain-mediated N-WASP activation. Our data not only provide novel mechanistic insight into the regulation of N-WASP at epithelial TJ (see below) but may also have important implications for other processes that are dependent on N-WASP, including invadopodia and lamellipodia formation, endocytosis, and actin-based motility of intracellular pathogens.

### ARHGAP12 suppresses N-WASP to dampen junctional tension and to promote TJ leak pathway permeability

We demonstrate that cells lacking ARHGAP12 exhibit increased junctional tension and reduced permeability to 4 kDa dextran. Although TJ formation was delayed in ARHGAP12 KO cells, ion permeability (i.e. the TJ pore pathway) was normal once the cell monolayer had matured. This suggests a specific function for ARHGAP12 in the control of the TJ leak pathway and strongly supports the notion that the pore and leak pathways are controlled independently of each other (Monaco et al., 2021; Shen et al., 2011). Since we have thus far not observed changes in the permeability to larger macromolecules (10 KDa dextran; data not shown), we propose that ARHGAP12 selectively controls the paracellular passage of small macromolecules up to 4 kDa in size. Importantly, both the tightening of the TJ permeability barrier and the increase in junctional tension observed in ARHGAP12 KO cells were restored upon Arp2/3 or myosin-II inhibition. Moreover, and importantly, reducing N-WASP levels in ARHGAP12 KO cells resulted in dextran flux rates comparable to those observed in wild-type cells. Together with our in vitro data this indicates that ARHGAP12 suppresses N-WASP to dampen Arp2/3-mediated branched actin assembly at TJ. This ultimately reduces the pool of TJ-associated F-actin that can be remodelled into actomyosin fibres, limiting junctional tension and promoting the flux of small macromolecules across the TJ (Figure 9).

It is well established that Arp2/3-mediated assembly of branched actin networks generates a pushing force against the membrane, which increases membrane tension (Papalazarou and Machesky, 2021). At adherens junctions, such pushing forces are critical for maintaining cadherin-based adhesions, which are otherwise pulled apart by myosin-II-mediated contractility (Efimova and Svitkina, 2018; Li et al., 2020). Although it is unclear at present whether an analogous push/pull system exists at the level of the TJ, it has been shown that myosin-II-mediated contractility regulates TJ mechanics and promotes leak pathway flux (Citi, 2019; Horowitz et al., 2023; Rouaud et al., 2023; Shen et al., 2006). However, too much tension is clearly detrimental to the TJ barrier (Choi et al., 2016; Fanning et al., 2012; Nguyen et al., 2024; Otani et al., 2019), suggesting that tension needs to be finely balanced to prevent barrier breakdown. There is also good evidence from work in cultured cells (Kovacs et al., 2011; Van Itallie et al., 2015; Verma et al., 2012; Wu et al., 2014) and from KO studies in mice (Garber et al., 2018; Kalailingam et al., 2017; Zhou et al., 2013) that N-WASP and the Arp2/3 complex promote F-actin assembly and the stability of F-actin networks to enhance junctional tension and to tighten the TJ permeability barrier. In addition, and interestingly, a dynamic interaction between ZO-1 and F-actin controls the paracellular transport of macromolecules without necessarily affecting ion transport (Belardi et al., 2020; Fanning et al., 2012; Spadaro et al., 2017; Van Itallie et al., 2009). Altogether the data indicate that N-WASP and Arp2/3-driven branched actin assembly strengthens the TJ barrier, possibly by generating an optimal level of tension that stabilizes the TJ strand network. We propose that ARHGAP12 is a negative regulator of this system that locally destabilizes TJ to promote leak pathway flux. Local and short-lived TJ leaks have indeed been observed during epithelial tissue shape changes in cultured cells and in Xenopus epithelia (Chumki et al., 2022; Richter et al., 2022; Stephenson et al., 2019; Varadarajan et al., 2022; Varadarajan et al., 2019). Interestingly, such TJ breaks are quickly repaired through F-actin assembly and myosin-II-dependent contractility (Stephenson et al., 2019; Varadarajan et al., 2022). It will be interesting to address in future whether N-WASP and Arp2/3 complex are required for the repair of TJ leaks and whether ARHGAP12 increases the frequency of TJ breaks or slows down the repair mechanism.

We also identified an interaction between ARHGAP12 and the WRC. Although both WAVE2 and Abi1 interacted with ARHGAP12 in IPs, the purified WAVE2 protein did not bind to the purified tWW domain in pulldown experiments. Since the WAVE2 PRD does not contain PPxR repeats, the interaction with the WRC may be mediated via an unknown intermediary protein. Regardless of the mode of interaction, the WRC is unlikely to be involved in the TJ permeability phenotype we observe in ARHGAP12 KO cells. Firstly, depletion of N-WASP in ARHGAP12 KO cells was sufficient to fully restore dextran flux, clearly demonstrating that N-WASP functions downstream of ARHGAP12 to control leak pathway permeability. Secondly, WAVE2 regulates F-actin assembly and junctional tension at adherens junctions rather than TJ (Verma et al., 2012). Indeed, all WRC components localise at a more basal junctional position than ARHGAP12 and N-WASP (Figure S1). We speculate, therefore, that the interaction with the WRC may be relevant in a different context, for example during the initial stages of cell-cell junction assembly. These ideas should be pursued in future.

In conclusion, our data indicate that ARHGAP12 acts as a molecular rheostat that fine-tunes F-actin assembly and tension at TJ to regulate paracellular transport of small macromolecules via the TJ leak pathway. We propose that ARHGAP12 functions via a two-tiered mechanism: a) it dampens SH3 domain-mediated N-WASP activation via a direct inhibitory interaction with the N-WASP PRD, and b) it locally suppresses Rac/Cdc42-GTP loading via its GAP activity, preventing GTPase-dependent activation of N-WASP (Figure 9). The work presented here provides novel mechanistic insight into how junctional F-actin assembly and tension controls the TJ permeability barrier and establishes a critical and specific role for ARHGAP12 in promoting leak pathway flux.

### Limitations of the study

We have analysed the functions of ARHGAP12 in MDCK cells, which is a commonly used model cell line to study epithelial TJ formation and permeability. Whether the mechanisms we describe in this cell culture model apply to other epithelial cells and epithelial tissues remains to be explored. In this study we focused on the function of the ARHGAP12 tWW domain in regulating the activity of N-WASP. To what extent the ARHGAP12 GAP activity contributes to the phenotypes we observe in ARHGAP12 KO cells needs to be addressed in future studies.

## Acknowledgements

We thank Walter Hunziker, Koh Cheng Gee, and Thirumaran Thanabalu for providing reagents and for valuable discussions, El Sahili Abbas for help with ITC measurements, and Bono Keir Audrey Mendoza, Goh Wei Xuan, Xinyi Lee, Tommy Lee Lam, and Zhiyi Lu for their contributions to this work. This work was supported by a Ministry of Education (MOE) Academic Research Fund Tier 2 grant (MOE-T2EP30121-0019) to AL.

## Author contributions

HMT generated MDCK cell lines and performed and analysed all MDCK cell-based experiments, YL cloned, expressed and purified recombinant proteins and performed and analysed all in vitro protein-protein interaction assays and COS7/293T cell transfections, KZ performed and analysed actin polymerisation and N-WASP oligomerisation assays, XT performed laser ablation experiments, YT supervised XT, YM supervised KZ, AL conceived the project, supervised HMT and YL, and wrote the manuscript.

## Conflict of interest

The authors declare no conflict of interest

## Materials and Methods

### DNA constructs and mutagenesis

ARHGAP12 cDNA was purchased from Source BioScience (clone ID #30528830). To create EGFP-ARHGAP12 and its variants, full-length ARHGAP12 cDNA and ARHGAP12 fragments were PCR amplified and inserted into pEGFP-C1 (Addgene #6084-1) using appropriate restriction sites. mCherry-ARHGAP12 was created by inserting the PCR amplicon into pmCherry-C1 (Clontech (TaKaRa) #632524). N-WASP cDNA was purchased from Addgene (plasmid #33019), PCR amplified and ligated into pEGFP-C1 plasmid using *XhoI* and *EcoRI* restrictions sites. For protein purification, the ARHGAP12 tandem WW (tWW) and SH3 domains, and the TUBA (GenBank #BC041628.1) and CIP4 (GenBank #BC013002.2) SH3 domains were cloned into a pET28b(+) vector, which places the 6×His tag and a thrombin cleavage site at the N-terminus of the insert. Point mutations in the tWW domain were introduced using site-directed mutagenesis. To produce proline-rich peptides (12-15 amino acids in length) fused to an N-terminal GST tag, complementary oligonucleotides were cloned into pGEX-6P-1 vector. The partial PRD of N-WASP (amino acids 261-332) was synthesized as a gBlock^TM^ gene fragment (Integrated DNA Technologies) and ligated into pGEX-6P-1 using *BamHI* and *EcoRI* restriction sites. Guide RNAs (gRNAs) targeting canine ARHGAP12 were ligated as complementary oligonucleotides into the PX459 CRISPR-Cas9 vector. All primers, oligonucleotides, and vectors can be found in Supplementary File 1.

### Protein Purification

The 6×His tagged ARHGAP12 tWW domain construct (His_6_-tWW) was transformed into *E. coli* BL21(DE3) competent cells. The bacteria were grown at 37°C, shaking at 200 rpm until the absorbance at 600 nm (OD600) reached ∼0.6. Protein expression was induced with 100 μM isopropyl-thio-β-D-galactoside (IPTG) at 16°C overnight at 160 rpm. The bacterial pellet was resuspended in Buffer A (25 mM HEPES pH7.0, 250 mM NaCl, 5 mM β-mercaptoethanol, 5% glycerol, protease inhibitor cocktail, 10 mM imidazole) and lysed by sonication. The cell suspension was centrifuged at 40,000 rpm for 20 min to pellet the debris. The supernatant was filtered through a 0.45 µm filter and injected into a HisTrap HP column (GE Healthcare) through an AKTA purifier (GE Healthcare). Samples were washed with Buffer B (25 mM HEPES pH7.0, 250 mM NaCl, 5 mM β-mercaptoethanol, 5% glycerol, 20mM imidazole) and eluted with increasing concentrations of imidazole in Buffer B. Fractions containing the tWW domain were concentrated using a Vivaspin turbo 10K MWCO (Satorius) spin column and passed through a HiLoad 16/60 Superdex 200pg SEC column (Cytiva) in SEC buffer (25 mM HEPES pH7.0, 250 mM NaCl, 1 mM Dithiothreitol (DTT), 5% glycerol). Peak fractions containing intact tWW domain were concentrated, flash frozen in liquid nitrogen and stored at -80°C. Similar protocols were used to express and purify the ARHGAP12 tWW mutants and the SH3 domains of ARHGAP12, TUBA, and CIP4.

GST-N-WASP-PRD and GST-peptide fusion constructs were cloned into pGEX-6P-1 vector, which places the GST-tag and PreScission site at the N-terminus. The sequenced vector was transformed into *E. coli* BL21(DE3) cells. Protein expression was performed at 16°C overnight via induction with 100 μM IPTG. Cells were harvested by centrifugation and lysed by sonication in binding buffer (25 mM HEPES, pH 7.4, 150 mM NaCl, 1 mM DTT, protease inhibitor cocktail (Thermo Scientific #A32965). The cell suspension was cleared by centrifugation at 40,000 rpm for 20 min. The supernatant was filtered through a 0.45 µm filter. The lysate was incubated with Glutathione Sepharose 4B (Cytiva) and washed with 10 column volumes of binding buffer using gravity flow columns. The GST tag protein was eluted in 50 mM HEPES, pH 8.0, 150 mM NaCl, 1 mM DTT, 20 mM reduced glutathione as 0.5 ml per fraction. Purity was verified by SDS-PAGE and fractions were concentrated using a Vivaspin turbo 10K MWCO (Satorius), flash frozen in liquid nitrogen and stored at -80°C. To purify GST-PRD, concentrated elutions were passed through a HiLoad 16/60 Superdex 200pg SEC column (Cytiva) in SEC buffer (25 mM HEPES pH7.0, 150 mM NaCl, 1 mM DTT, 5% glycerol). The fractions containing full-length intact GST-PRD protein were concentrated, flash frozen in liquid nitrogen, and stored at -80°C.

GST-SH3-tWW, GST-SH3 and GST-tWW were transformed into *E. coli* BL21(DE3). Protein expression was induced with 200 μM IPTG. Bacterial cultures were grown at 37°C for 4 hours or 16°C overnight at 160 rpm. Cells were resuspended in lysis buffer (PBS, pH 7.4, 1 mM DTT, 1 mg/ml lysozyme, 150 mM NaCl and protease inhibitor cocktail and lysed with LM20 Microfluidizer (Microfluidics™) at 20,000 Pa three times. The lysate was incubated with Glutathione Sepharose 4B beads (Cytiva) for 2 hours at 4°C on BenchRocker™ 2D (Benchmark). Flow through (FT) was collected and washed 3 times with ice-cold PBS. Proteins were eluted five times from the beads with elution buffer (50 mM Tris-Cl pH 8.0, 150 mM NaCl, 1 mM DTT, 20 mM reduced glutathione, 25% glycerol). Elution factions were pooled and dialyzed overnight against dialysis buffer (50 mM Tris-Cl pH 8.0, 150 mM NaCl, 1 mM DTT, 25% glycerol).

6×His tagged tri-SH3 (derived from FBP17) and 6×His tagged full-length N-WASP were transformed into BL21(DE3) Rosetta cells. A 50 ml overnight culture was inoculated into 2 liters of TB medium. The cells were then grown at 37°C until they reached an OD600 of ∼0.6. Protein expression was induced with 100 µM IPTG at 16°C for 16 hours. Cells were harvested by centrifugation and resuspended in 45 ml of Binding Buffer (20 mM HEPES, pH 7.4, 500 mM NaCl, 20 mM Imidazole). Cell disruption was achieved using a homogenizer (LM20 Microfluidizer™) in Binding Buffer supplemented with 1 mM PMSF, 0.1% (v/v) Triton X100, and protease inhibitor cocktail. The supernatant was obtained by centrifugation at 20,000 rpm for 1.5 hours at 4°C, filtered (0.22 μm), and then applied onto a 5 ml HisTrap HP column (G.E. Healthcare) connected to an FPLC AKTA system (G.E. Healthcare). The column was washed with Washing Buffer (20 mM HEPES, pH 7.4, 50 mM imidazole, and 500 mM NaCl), and proteins were eluted with Elution Buffer (20 mM HEPES, pH 7.4, 500 mM imidazole, and 500 mM NaCl). The protein was further purified by size exclusion chromatography on a HiLoad 16/600 Superdex 200pg column (G.E. Healthcare) in Gel filtration Buffer (20 mM HEPES pH 7.4, 500 mM NaCl, 10% Glycerol, and 1 mM DTT), and finally concentrated to a concentration of ∼5 mg/ml using 15 ml 50 kDa cut-off concentrators (Amicon Inc.).

To produce fluorescently labeled N-WASP for single particle imaging, the primary amine group of N-WASP was labeled using an Alexa Fluor™ 488 labeling kit (Thermo Scientific). Briefly, the protein of interest was prepared at ∼2 mg/ml in a buffer containing 0.1 M sodium bicarbonate buffer. The Alexa dye was added to the mixture and incubated overnight at 4°C. Excess dye was removed by using a 5 ml HiTrap Desalting column (G.E. Healthcare) in gel filtration Buffer (20 mM HEPES pH 7.4, 500 mM NaCl, 10% Glycerol, and 1 mM DTT). Labeling efficiency was determined using a Nanodrop 2000 (Thermo Scientific).

### Immunoprecipitation (IPs)

For IPs, 293T or MDCK-II cells were lysed in ice-cold IP buffer (IPB) (25 mM Tris pH 8.0, 150 mM NaCl, 5 mM ethylenediaminetetraacetic acid (EDTA), 0.5% Triton X-100, and protease inhibitor cocktail (ThermoFisher #A32965) for 15 min. Lysates were cleared by centrifugation at 15,000 g for 30 min at 4℃. IPs of GFP-tagged proteins were performed by incubating cleared cell lysates for 2 hours at 4°C with GFP-Trap® beads (ChromoTek) that had been pre-equilibrated with cold IPB. Following incubation, beads were pelleted via centrifugation at 500 g for 2 min at 4°C. Bound proteins were eluted with SDS-PAGE loading dye and boiled at 95 ℃ for 2-5 min. The samples were analysed by Western blotting.

### GST pulldown assay

For pulldowns with MDCK cell lysates 20 μg of purified GST, GST-SH3-tWW, GST-SH3 and GST-tWW were coupled to 10 μl of Glutathione Sepharose 4B beads (GE Healthcare Life Sciences) in 450 μl of GST pulldown buffer (50 mM HEPES pH7.4, 100 mM NaCl, 5% glycerol, 0.1% Tween 20, 1% BSA) for 1.5 hours at 4℃. The supernatant was discarded and the beads were washed three times with PBS. Beads were then incubated with 400 μg of MDCK-II cell lysates (in a total volume of 400 µl) for 1.5 hours at 4°C and washed three times with PBS. Bound proteins were eluted from the beads by boiling for 5 min in SDS-PAGE loading dye. For pulldowns with two purified proteins 20 μg of purified GST, GST-N-WASP-PRD or GST-tagged peptides were coupled to 10 μl of Glutathione Sepharose 4B beads (see Supplementary File 1 for the list of GST-peptide sequences). After several washes, equimolar amounts of purified His-tag fusion protein in 450 μl of GST pulldown buffer was added to the beads. The mixture was rotated in the cold room for 1h. The beads were washed four times with GST pulldown buffer (without BSA). Bound proteins were eluted with SDS-PAGE loading dye and boiled at 95 ℃ for 2-5 min. The samples were analysed by Western blotting. The GST pulldown competition assays followed a similar protocol. After coupling the beads with the GST-PRD or GST protein, five times molar excess of His-tag protein (over GST-PRD) was added to saturate PRD binding. Binding was performed in the cold room for 1h. The beads were washed three times with GST pulldown buffer, followed by the addition of the second His-tag protein at the indicated molar ratios (His-tag protein over GST-PRD). The beads were incubated rotating in the cold room for another 1h. The beads were washed four times with GST pulldown buffer (without BSA), eluted with SDS-PAGE loading dye, and boiled at 95 ℃ for 2-5 min. Bound proteins were analysed by Western blotting.

### SDS-PAGE and Western Blotting

Samples were loaded into the wells of 1.0 mm poly-acrylamide gels of the appropriate gel percentage and ran at 100-120V in SDS running buffer (25 mM Tris, 190 mM glycine, and 0.1% SDS). Wet transfer setup was performed at 70V for 3 hours or 20V overnight in transfer buffer (25 mM Tris and 190 mM glycine) to transfer the protein onto a methanol-activated Polyvinylidene difluoride (PVDF) membrane. The PVDF membranes were dehydrated in methanol for 10 sec and dried for at least 1 hour. Dried membranes were incubated with primary antibodies (see list of antibodies in Supplementary File 1) diluted in PBST (PBS, 0.2% Tween-20) with 2% bovine serum albumin (BSA) at room temperature for 1 hour or 4°C overnight. Membranes were washed three times for 5 min each with PBST before incubation with horseradish peroxidase (HRP)-conjugated secondary antibodies diluted in PBST with 5% fat-free milk. Membranes were further washed with PBST three times and developed with a chemiluminescent substrate (Merck).

### Coomassie Staining

Gels were fixed with hot Coomassie Blue staining solution (40% H_2_O, 50% methanol, 10% acetic acid (v/v/v), 0.25% Coomassie Brilliant Blue G250) for 10 min. Gels were then destained with a destaining solution (40% H_2_O, 50% methanol, 10% acetic acid (v/v/v)) until the desired background was achieved.

### Isothermal Titration Calorimetry (ITC)

ITC experiments were conducted on a Microcal ITC200 (Malvern Pananalytical) at 20 °C. All peptides were synthesized and purified by Genscript (purity: 95%, all peptide sequences are listed in Supplementary File 1). All protein samples were cleared by centrifugation before the experiment. Titration buffer contained 25 mM HEPES (pH 7.4) and 150 mM NaCl. Each titration point was performed by injecting 1 µl of peptide ligand sample (500 µM) into the cell containing target protein (50 µM) within 2 sec at a time interval of 150 sec to ensure that the titration curve returned to the baseline. 20 injections were performed to saturate the binding events. The titration peaks and measurements were analysed by the MicroCal PEAQ-ITC Analysis Software (Malvern Pananalytical). All measurements were repeated two to three times with buffer control experiments.

### *In vitro* actin assembly assay

The *in vitro* real-time actin assembly assays were performed on Biotin-PEG coated glass slices (Laysan Bio Inc) sectioned by 6-well chamber slices (Ibidi). The chamber was first blocked by 30 µl HBSA buffer (20 mM HEPES pH7.4, 1 mM EDTA, 50 mM KCl, 1%(m/v) BSA) and incubated for 30 sec. The glass surface was then conjugated with streptavidin by adding 30 µl HEKG10 buffer (20 mM HEPES pH7.4, 1 mM EDTA, 50 mM KCl,10%(v/v) glycerol, plus 0.1 mg/ml streptavidin) and incubating for 1 min. Afterward, 1x TIRF buffer (10 mM imidazole pH 7.4, 50 mM KCl, 1 mM MgCl_2_, 1 mM EGTA, 50 mM DTT, 0.3 mM ATP, 20 nM CaCl_2_, 15 mM glucose, 100 mg/ml glucose oxidase, 15 mg/ml catalase, 0.25% methylcellulose) was used to wash away free streptavidin. Next, 30 μl 3X protein-actin mix containing 3 μM G-actin (89% purified globular rabbit actin, 10% Oregon Green 488-actin, 1% Biotin-actin), 200 mM EGTA, 110 mM MgCl_2_, and desired protein solution was mixed with 30 μl 2X TIRF buffer and added into the chamber to a final volume of 90 μl to initiate the actin polymerization. Image stacks with 15 sec intervals for 15min were acquired at room temperature using a Nikon Ti2-E inverted microscope equipped with a 100x 1.45NA Plan-Apo objective lens and a TIRF module (iLasV2 Ring TIRF, GATACA Systems) and an ORCA-Fusion sCMOS camera (Hamamatsu Photonics). Imaging laser was provided 488 nm/150 mW (Vortran) in a laser launch (iLaunch, GATACA Systems). Focus was maintained by hardware autofocus (Perfect Focus System), and image acquisition was controlled by MetaMorph software (Molecular Device). To quantify the signal of actin filaments, 22x22 μm^2^ ROIs were chosen from each time point image, N=9 and 18 for actin control or without protein, respectively.

### Quantification of Single Particle Intensity

The protein mixture of 20 nM AF488-N-WASP and the desired WW concentration was premixed in PBS buffer in a final volume of 200 µl. After 15 min of incubation, the protein mixture was added and in a 96-well glass bottom plate (Cellvis). TIRFM imaging was then performed immediately with consistent parameters. To quantify the intensity of AF488-N-WASP particles, we analyzed the images using Trackmate in ImageJ. First, a preliminary particle analysis was performed manually, setting the estimated diameter for particle detection to 0.65 μm (10 pixels). In addition, quality control was performed to filter out false selections from the background and ensure that only visible punctate protein signals were selected. The selected particles were then analyzed for signal intensity in Trackmate. Specifically, 200 from over 1000 particles of each protein combination were quantified from at least ten images to ensure statistical robustness.

### Mammalian cell transfection

Cos-7 (African green monkey kidney fibroblast-like) cells, HEK293T cells and Madin Darby Canine Kidney type II (MDCK-II) cells were grown in DMEM++ (DMEM (4.5 g/L glucose) containing 10% fetal bovine serum (FBS) and 100 U/ml penicillin-streptomycin) at 37°C with controlled CO_2_ level at 5%. Cells were transfected with expression plasmids using Lipofectamine-3000 according to the manufacturer’s protocol (Invitrogen, cat.no. L3000008).

### Generation of CRISPR/Cas9 KO Cells

The design of gRNAs was facilitated through the CRISPR Direct webpage (http://crispor.tefor.net/) and were purchased as short oligonucleotides and inserted into the vector using BbsI-HF® (NEB) restriction enzyme. gRNAs were co-transfected with pEGFP-C1 vector in a ratio of 4:1 using Lipofectamine-3000. Following transfection, cells underwent a selection process with 500 µg/ml G418 for 2 weeks before being subjected to FACS single-cell sorting to isolate GFP-expressing cells. Cells were subjected to various assays to confirm the knockout of ARHGAP12 in the obtained clones. This included procedures such as western blotting, genomic DNA extraction, and immunofluorescence. ZO-1 and ZO-2 CRISPR KO cells were a kind gift from Walter Hunziker, IMCB, AStar, Singapore.

### Generation of shRNA Lines

Short hairpin RNAs (shRNAs) were synthesized as complementary oligonucleotides by BioBasic Asia, Singapore. The oligonucleotides were designed with BglIII (5’) and HindIII (3’) sticky ends and annealed in annealing buffer (10 mM Tris pH 7.5, 50 mM NaCl, 10 mM EDTA). The annealed shRNAs were then ligated into the pSuper vector (Oligogene) and verified by Sanger sequencing, which included 5% dimethyl sulfoxide (DMSO) in the reaction mix to facilitate hairpin melting. The shRNAs were transfected into MDCKII cells using Lipofectamine-3000 at a LF3000/DNA ratio of 3:1. To generate stable shRNA cell lines, the cells were cultured under selection for 2 weeks in DMEM containing 200 µg/ml hygromycin B. Western blotting was performed to confirm the efficiency of the shRNAs.

### Calcium Switch Assay

MDCK-II cells were seeded in 24-well plates containing glass coverslips and grown in DMEM++ until they reached 80% confluency. Upon reaching confluency, DMEM++ media was switched to low calcium DMEM (5% FBS, 1x Glutamax, 1 mM sodium pyruvate, 100 U/ml PS, and 3 µM calcium chloride (CaCl_2_)). Cells were incubated for 16-24 hours before switching back to DMEM++ to allow junction formation. Cells were fixed at different time points and processed for immunofluorescence. Average junctional length quantification of the MDCKII WT and MDCKII ARHGAP12 KO cells was performed in FIJI (Image J) by measuring the length of ZO-1-positive cell junctions divided by the total number of cells.

### Immunofluorescence

MDCK-II cells, Cos-7 cells or HEK293T cells were fixed for immunofluorescence using 1% or 4% paraformaldehyde (PFA) or by methanol fixation, depending on the antibodies used. For cells fixed with PFA, cells were washed three times and then permeabilized with 0.1% TritonX-100 in PBS for 10 min. After three washes with PBS cells were blocked in 0.4 µm filtered PBS containing 10% FBS for 2h at room temperature or 4°C overnight. Samples were incubated with primary and secondary antibodies (see Supplementary File 1) diluted in PBS, 0.1% BSA, 0.01% Tween-20. F-actin staining was performed with Phalloidin-Atto 565 (Sigma cat.no. 94072). DNA was counterstained using DAPI (Sigma, cat.no. D9542). Cells were mounted in Vectashield antifade mounting medium (Vector Laboratories, cat.no. H-1000). Samples were imaged using a Spinning Disk Microscope (Corrsight, FEI Company) equipped with an Orca R2 CCD camera (Hamamatsu) using a 40X/1.3 oil or 63X/1.4 oil objective lens (Zeiss), utilizing 405, 488, 561 nm, and 633 nm lasers for excitation. High-resolution images were collected from Zeiss LSM 980 with Airyscan 2, with a 40 X/1.3 oil objective, utilizing 2% laser power of 405 (DAPI), 488 (GFP), and 561 nm (Red) lasers for excitation. The images were scanned 2788 × 2788 pixels, bi-directionally with scan zoom 2 or 4 and SR-4Y mode. Detection gain was within the range of 650 to 850.

### Time-lapse live cell imaging

Time-lapse live cell imaging was performed using a Corrsight spinning disk microscope (FEI Company) at 5% CO_2_ supply and 37°C. Cos-7 cells or HEK293T cells were seeded on fibronectin-coated 35 mm glass bottom dishes with a #1.5 coverslip (P35G-1.5-14-C, MatTek) and co-transfected with mCherry-LifeAct and EGFP-SH3-tWW or GFP-N-WASP and mCherry-ARHGAP12 using lipofectamine-3000 (see mammalian cell transfection section). Cells were imaged 20 hours post-transfection. Time lapse movies were acquired for up to ∼10 min using a frame rate of 10 to 30 sec. 488 (GFP) and 561 nm (Red) lasers were used for excitation using minimal laser power and exposure times. Movies were processed in FIJI (Image J).

### Transepithelial Electrical Resistance (TEER)

Cells were seeded onto Transwell® (Corning) filter inserts at a concentration of 2 × 10^5^ cells/cm^2^. The electrical resistance across the cell monolayer was monitored daily from the second day onwards for 7 days using the Millicell-ERS epithelial volt-ohm meter (Millipore). Transepithelial electrical resistance (TEER) was calculated by:

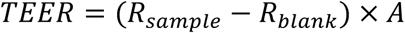

R_sample_ is the resistance of cell monolayer (Ohms), while R_blank_ is the resistance of the culture insert without cells (Ohms) and A is the surface area of the insert (cm^2^).

### Dextran Permeability Assay

Cells were seeded onto Transwell® (Corning) filter inserts at a concentration of 2 × 10^5^ cells per square centimeter for 8 days. 1 mg/ml of FITC-Dextran 4 kDa was placed onto the apical side of the Transwell insert. After 2, 4, and 6 hours, aliquots were taken out from the basolateral side for measurement and placed inside a 96-well Optical Coverglass Base Microplate (Thermofisher). Cells were either left untreated or treated with 50 µM Blebbistatin (Sigma-Aldrich), 200 µM CK-666 (MedChemExpress), and 10 µM SMIFH2 (Sigma-Aldrich). All inhibitors are listed in Supplementary File 1. The samples were then excited at 488 nm and the emission was read at 518 nm using Tecan Spark 10M Microplate Reader. Apparent permeability (P_app_) was quantified by:

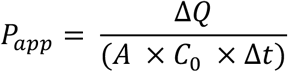

ΔQ is the amount of dextran detected on the basolateral side (mg), A represents the surface area of the insert (cm^2^), C_0_ is the initial concentration in the apical compartment (mg/ml), and Δt is the time of the experiment (s).

### Laser Ablation

Laser ablation experiments were conducted on MDCK-II WT and ARHGAP12 KO cells using a Nikon A1R MP confocal microscope equipped with an Apo 40× WI λ S DIC N2, N. A 1.25 objective. The cells were cultured to confluency on a MatTek dish and stained with CellMask™ (Invitrogen) to visualize the plasma membrane. Cells were subjected to either no treatment or treatment with 50 µM Blebbistatin (Sigma-Aldrich), 200 µM CK-666 (MedChemExpress), and 10 µM SMIFH2 (Sigma-Aldrich) for 4 hours. The laser employed for ablation had a wavelength of 355 nm, a pulse duration of 300 ps, and a repetition rate of 1 kHz. A PowerChip PNV-0150-100 from teeming photonics was utilized, and the laser power at the back aperture of the objective was set to 100 nW with a duration of 2 sec. The movement of tri-cellular junctions during laser ablation was monitored using the PointPicker plug-in in ImageJ. Recoil velocity was calculated using these equations:

1. Linear fitting equation – to calculate the velocity before laser ablation:

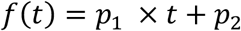 *p*_1_ is the slope of the linear equation while *p*_2_ is the rate at which junctional length changes occur, while t is the time (s)
2. Double exponential equation – to fit and calculate the initial recoil velocity:

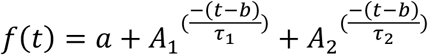 A, A1, A2, b, *τ*_1_ and *τ*_2_ are parameters to fit the equation. t is the time (s)
3. Double exponential equation – to fit and calculate cells with low velocity:

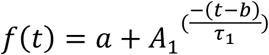 A, A1, and *τ*_1_ are parameters to fit the equation.

### Quantification and Statistical Analysis

Confocal images and time lapse movies were processed and analyzed in FIJI (ImageJ). The quantification of mean fluorescence intensity of N-WASP signal was done on maximum intensity projections of individual GFP positive or non-transfected cells. To quantify the co-localization of GFP-ARHGAP12 variants with ZO-1 or N-WASP, cells were outlined using the ‘Polygon Section’ tool and designated as regions of interest (ROIs) in the ‘ROI Manager’ tool in ImageJ. Pearson’s correlation coefficients were determined using the ‘Coloc2’ plugin in ImageJ. To quantify the fluorescence intensity of ARHGAP12 at cell-cell junctions, the junctions were outlined using the ‘Segmented Line’ tool of ImageJ. The ‘Measure’ tool was used to measure the fluorescence intensity inside and outside the box. Non-junctional fluorescence signal was subtracted. Unless stated otherwise, statistical analysis was performed using a two-tailed unpaired Student’s t-test in GraphPad Prism. Statistical significance was indicated as follows: n.s., not significant; * p ≤ 0.05; ** p ≤ 0.01; *** p ≤ 0.001; **** p ≤ 0.0001. Values are presented as mean ± SEM unless otherwise stated.

## Supplementary Movies

Movie S1: Live cell imaging of 293T cells co-transfected with GFP-N-WASP (left) and mCherry-ARHGAP12 (center). The panel on the right shows the merged channels.

Movie S2: Live cell imaging of COS7 cells co-transfected with LifeAct-mCherry (left) and GFP-SH3-tWW (center). The panel on the right shows the merged channels.

Movie S3: Representative laser ablation experiment of WT MDCK-II cells (left) and ARHGAP12 MDCK KO cells (right).

## Supplemental Figure Legends

**Figure S1:**
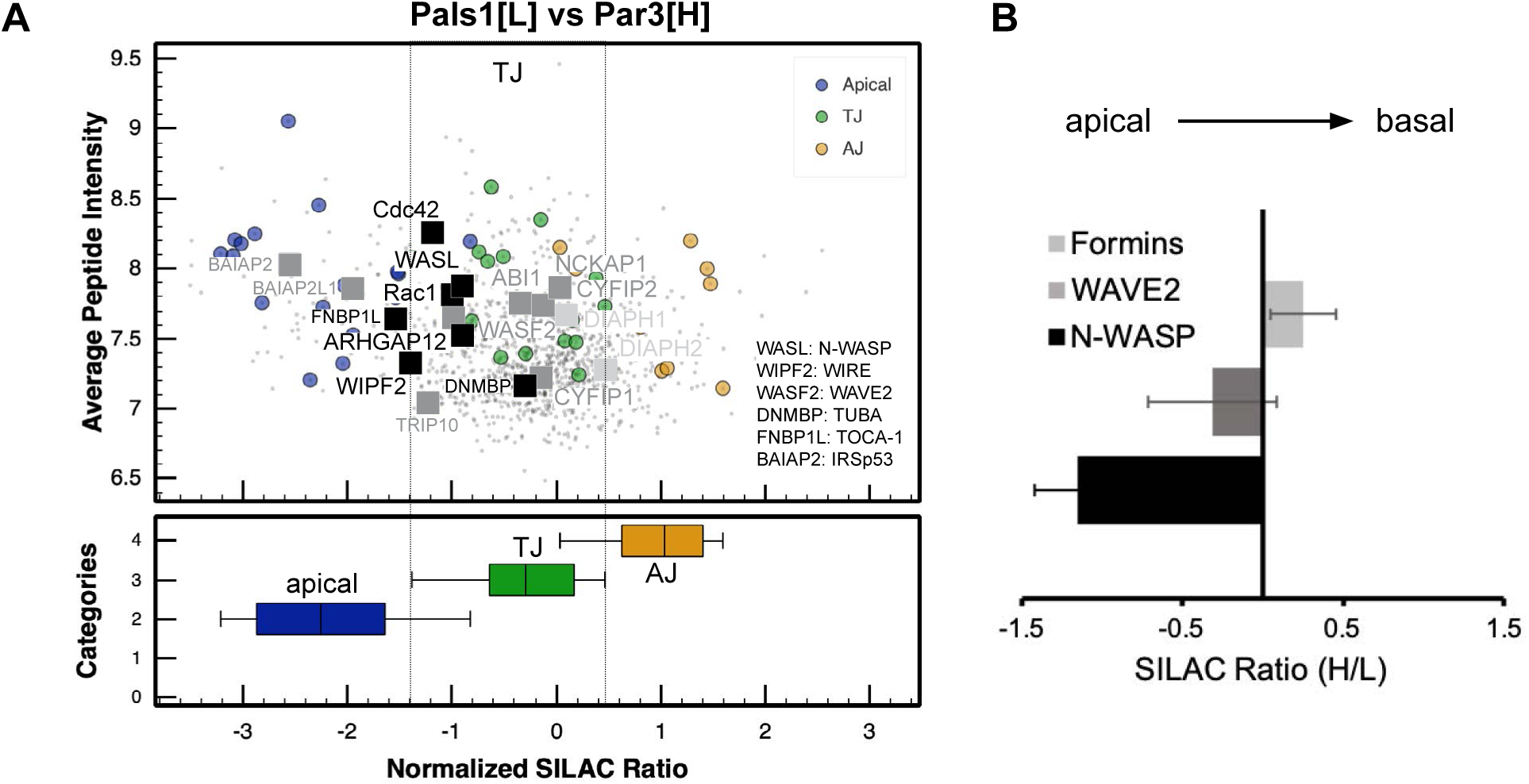
Sub-junctional localisation of ARHGAP12 and regulators of the actin cytoskeleton. (A) Scatter plot showing the sub-junctional localisation of ARHGAP12, N-WASP, WIRE, Rac1, Cdc42, the WRC (WAVE2, ABI1, ABI2, NCKAP1, CYFIP1, CYFIP2), formins (DIAPH1 and DIAPH2), and regulators of N-WASP and WAVE2 (e.g. TOCA-1, IRSp53, TUBA, TRIP10). This spatial map was determined by a ratiometric proximity proteomics approach using a Pals1/Par3 APEX2 SILAC pair (Tan et al., 2020). (B) Box plot showing the average apico-basal localisation of N-WASP-related proteins (N-WASP, WIRE, ARHGAP12, Rac1, Cdc42, TOCA-1), WAVE2-related proteins (the WRC), and formins (DIAPH1 and DIAPH2). Note that N-WASP and ARHGAP12 are localised at the apical aspect of the tight junction, whilst the WRC and formins are localised more basally.

**Figure S2:**
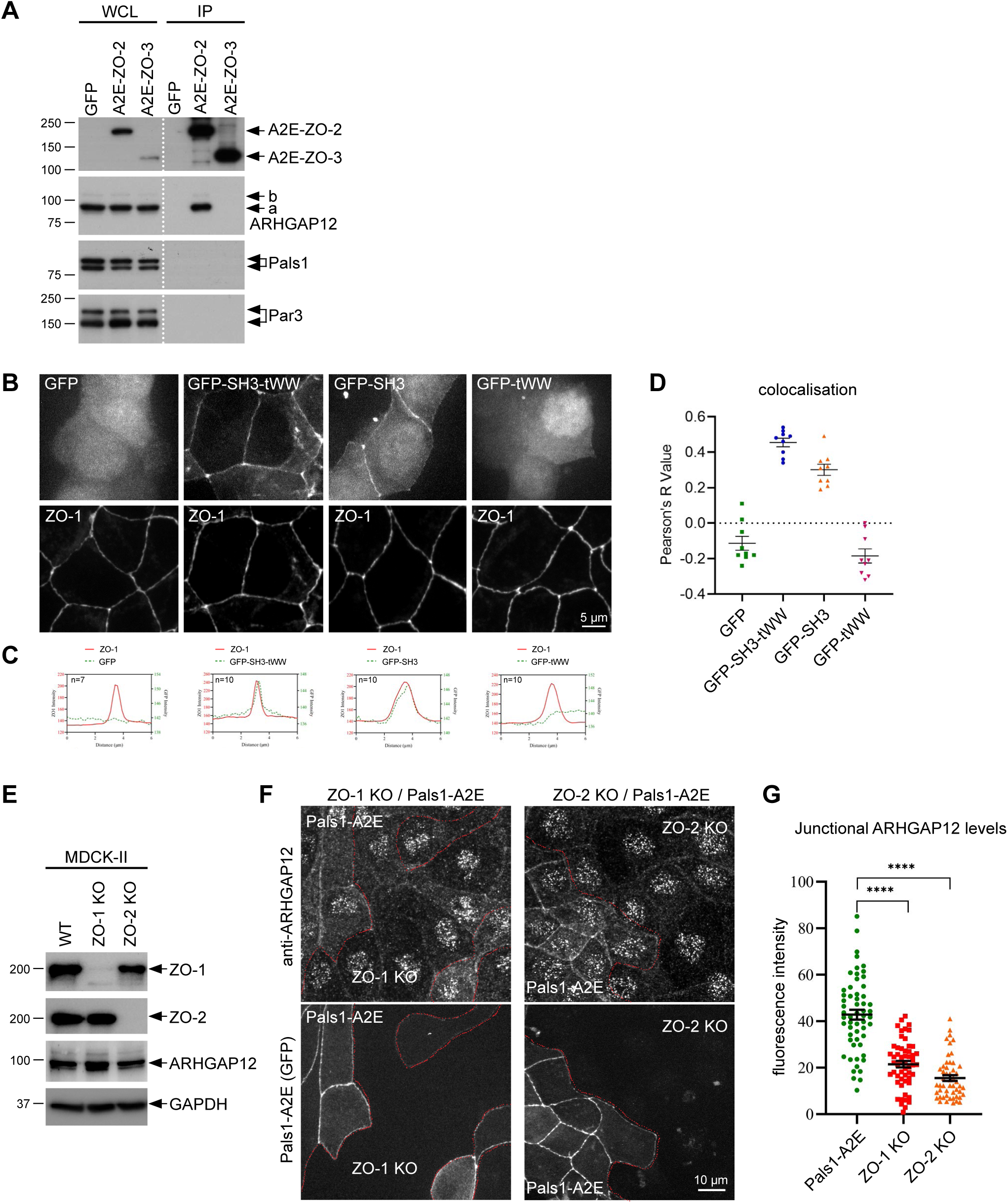
ARHGAP12 is recruited to tight junctions via ZO-1 and ZO-2. (A) Immunoprecipitation analysis of MDCK-II cells stably transfected with APEX2-EGFP fusion proteins of ZO-2 (A2E-ZO2), ZO-3 (A2E-ZO3), or EGFP as a control. GFP proteins were isolated using anti-GFP antibodies. (B) Immunofluorescence of MDCK-II cells transiently transfected with GFP, GFP-SH3-tWW, GFP-SH3, or GFP-tWW. Cells were stained with anti-ZO-1 antibodies. Scale bar 5 µm. (C) Line scan analysis of ZO-1 and GFP signal intensities of the data shown in (B). (D) Pearson’s Correlation (colocalization) analysis of the data shown in (B). (E) WB analysis of wild-type (WT), ZO-1 and ZO-2 KO MDCK-II cells. (F) Immunofluorescence of ZO-1 and ZO-2 KO MDCK-II cells co-cultured with MDCK-II wild-type cells stably transfected with Pals1-A2E. Cells were stained with anti-ARHGAP12 antibodies. (G) Quantification of junctional ARHGAP12 intensity in WT (Pals1-A2E), ZO-1 and ZO-2 KO MDCK cells. Error bars indicate SEM. **** = p<0.0001 (student’s t-test).

**Figure S3:**
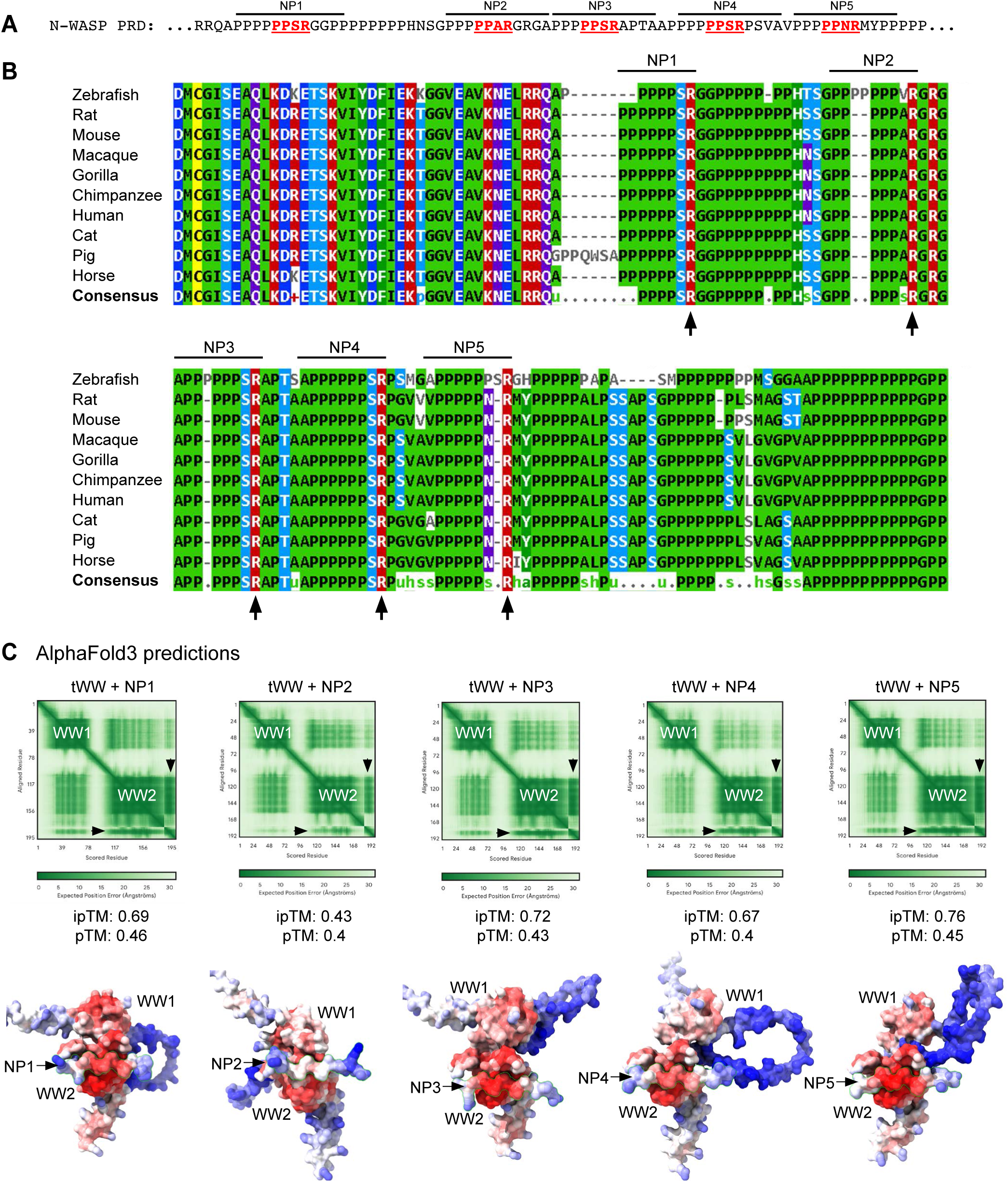
The PPxR repeats in the N-WASP PRD are highly conserved and are predicted to interact with the ARHGAP12 tWW domain. (A) Amino acid sequence of the human N-WASP PRD indicating the five conserved PPxR repeats (NP1, NP2, NP3, NP4, and NP5). (B) Multiple Sequence Alignment of the N-WASP PRD from various vertebrate species. The five PPxR repeats NP1-NP5 are indicated. (C) AlphaFold3 predictions of each PPxR peptide (NP1-NP5) with the tWW domain of ARHGAP12. Top: Predicted Aligned Error (PAE) plots, demonstrating that AlphaFold accurately predicts the WW1 and WW2 domain folds. The peptides NP1-NP5 are indicated by arrowheads. The ipTM and pTM values signify that the predictions are of medium to high confidence. Note that the tWW/NP2 prediction has relatively low confidence. Bottom: Surface representations in b-factor colour code. Red: high confidence, blue: low confidence.

**Figure S4:**
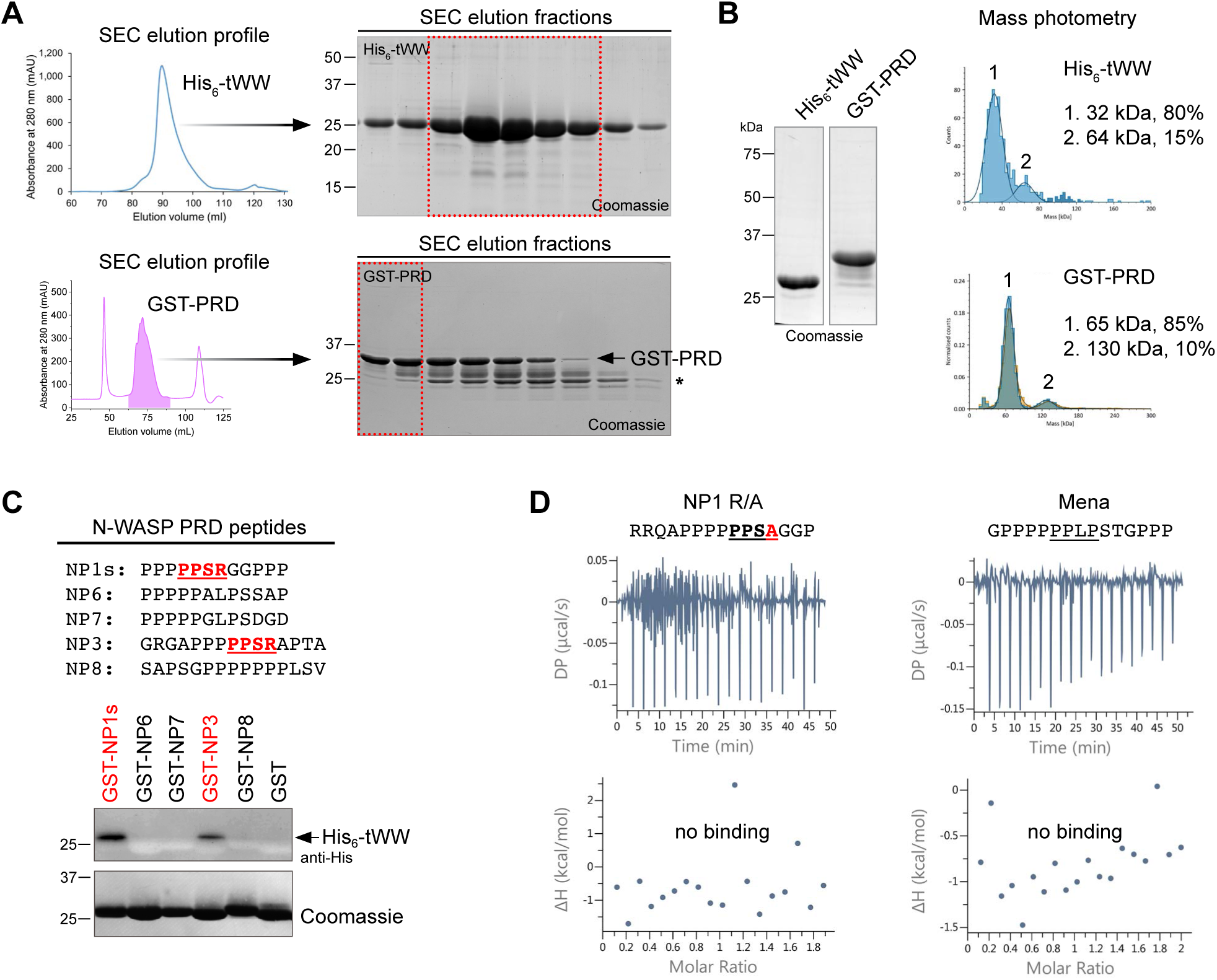
Purification and in vitro characterisation of the ARHGAP12 tWW domain. (A) Purification of His_6_-tWW and GST-PRD (of N-WASP) using affinity chromatography and Size Exclusion Chromatography (SEC). SEC elution profiles and Coomassie stained gels of the peak elution fractions are shown. (B) Coomassie-stained gel and mass photometry data of the purified His_6_-tWW and GST-PRD proteins. (C) The purified tWW domain binds specifically to PPxR motif containing peptides. GST fusion proteins of the indicated peptides were coupled to glutathione sepharose and incubated with His_6_-tWW. GST-PRD and GST was used as positive and negative control, respectively. (D) ITC data of the tWW domain and the mutant NP1 peptide (R/A mutation) or a Mena peptide containing PPLP motifs. No binding was observed in either case.

**Figure S5:**
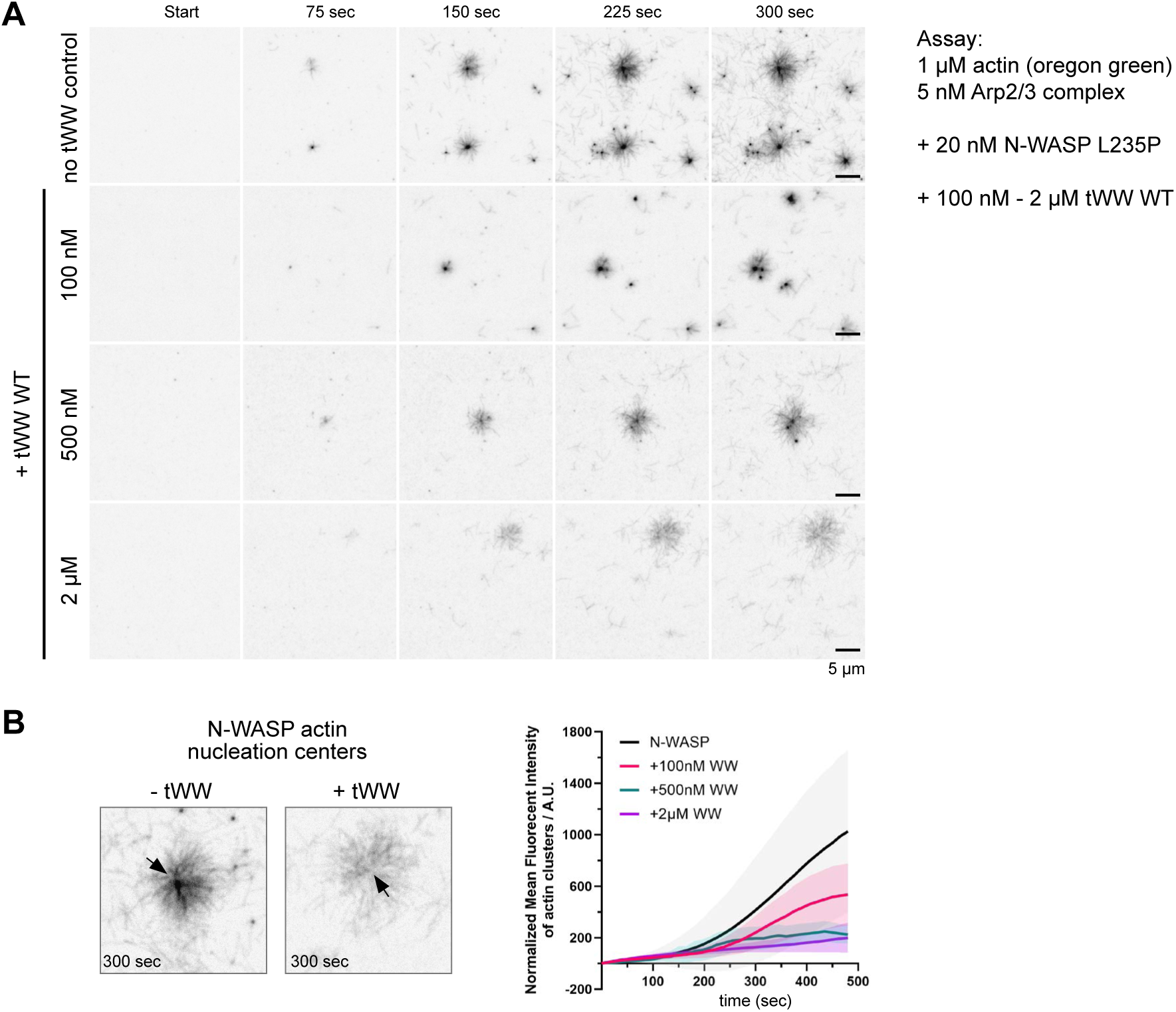
The ARHGAP12 tWW domain attenuates actin nucleation driven by a constitutively active N-WASP mutant. (A) In vitro actin polymerisation assay. Fluorescently labelled (Oregon green) G-actin, Arp2/3 complex, and the constitutively active N-WASP mutant L235P were incubated with increasing concentrations of the wild-type ARHGAP12 tWW domain. Actin polymerisation was monitored live using TIRF microscopy. (B) Quantification of the fluorescence intensity of N-WASP-induced F-actin clusters. Note that the tWW domain reduces the fluorescence intensity of F-actin nucleation centres (arrows) in a concentration-dependent manner.

**Figure S6:**
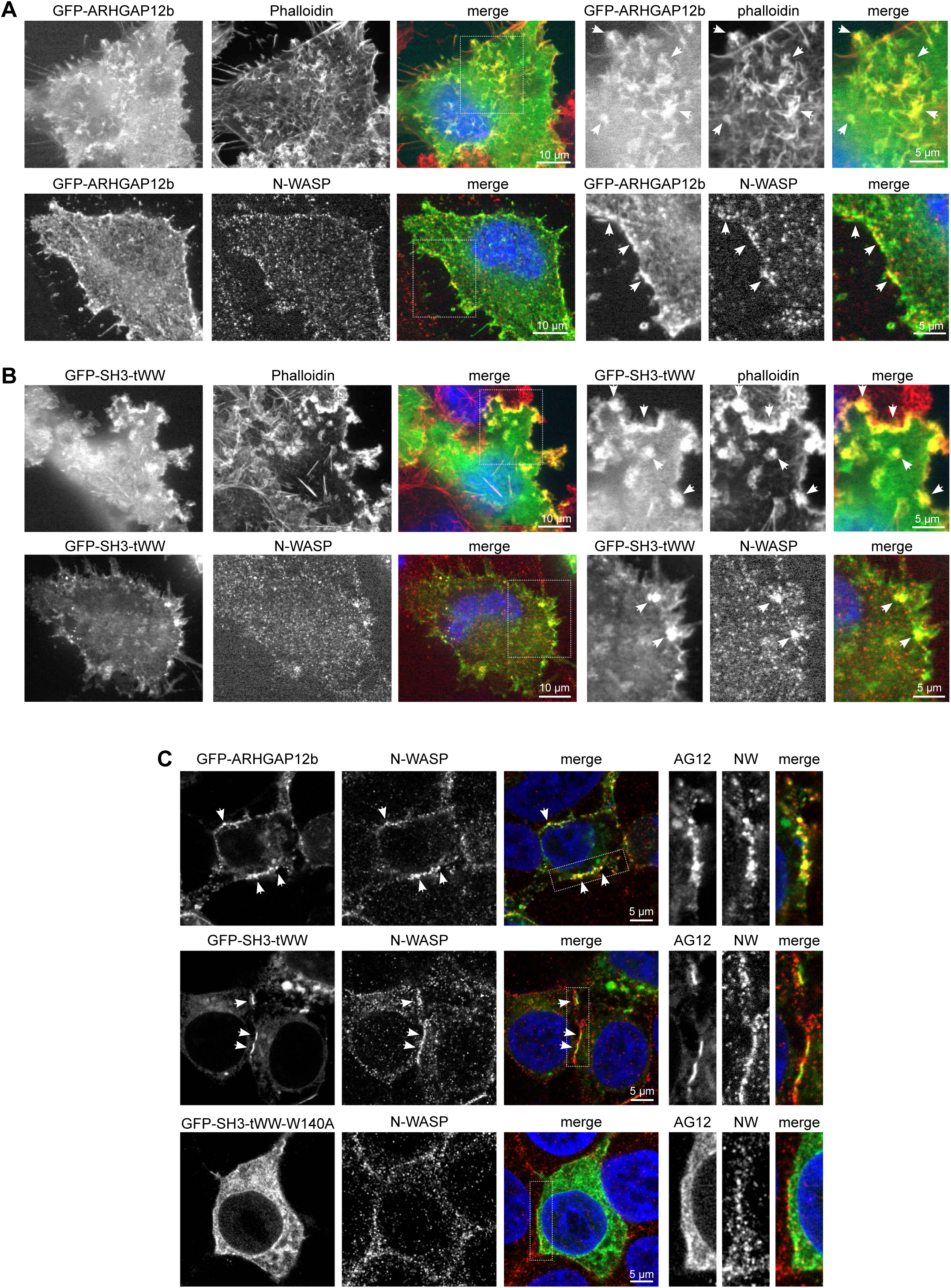
ARHGAP12 co-localises with F-actin and N-WASP. (A and B) COS7 cells transfected with GFP-ARHGAP12b (A) or GFP-SH3-tWW (B) were fixed, stained with Phalloidin to visualise actin or stained with anti-N-WASP antibodies, and imaged by confocal microscopy. Arrowheads highlight areas of colocalization. (C) 293T cells transfected with GFP-ARHGAP12b, GFP-SH3-tWW, or the GFP-SH3-tWW W140A mutant were fixed, stained with anti-N-WASP antibodies, and imaged by confocal microscopy. Arrowheads highlight areas of colocalization.

**Figure S7:**
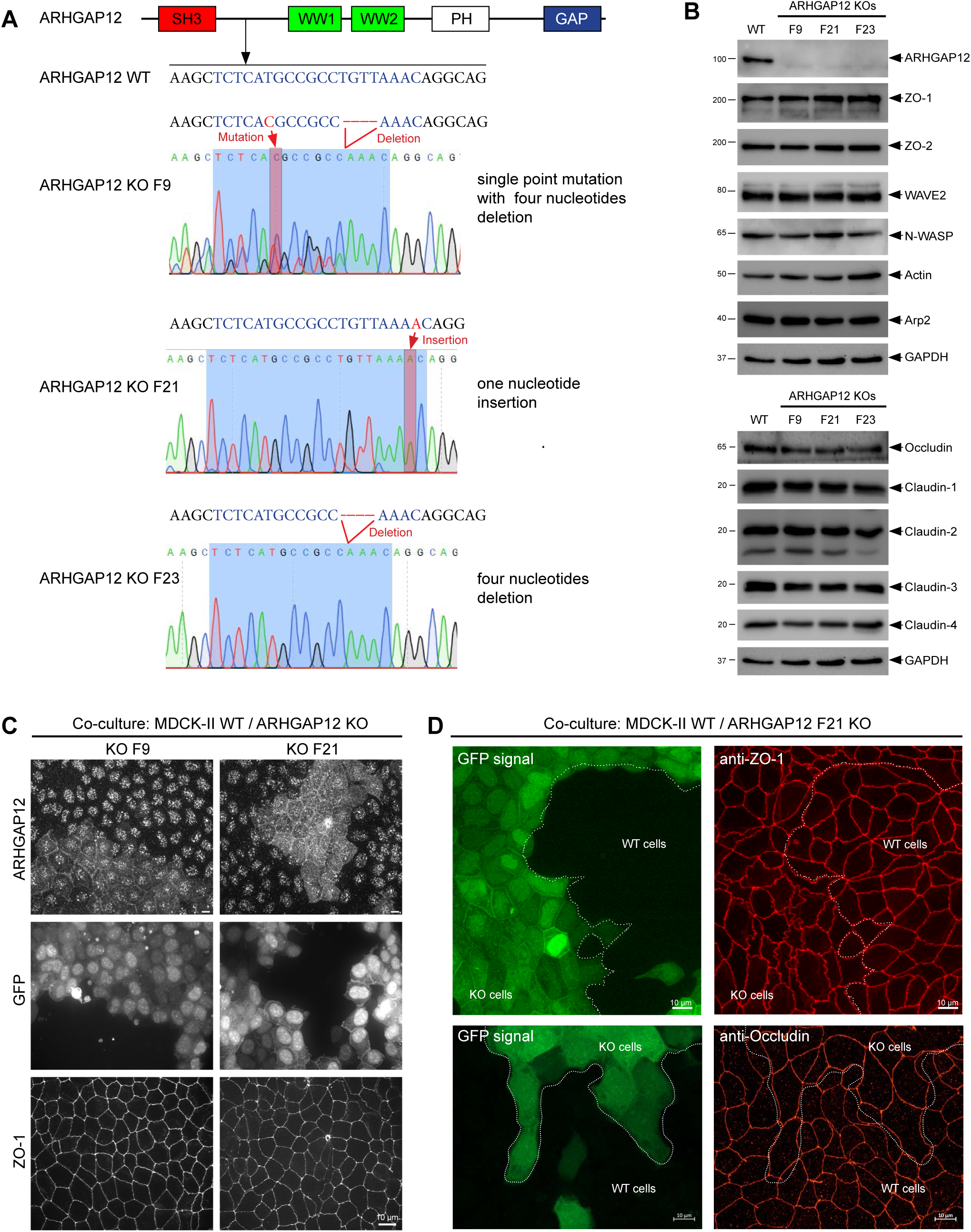
Generation and characterisation of ARHGAP12 MDCK-II KO cells. (A) Genomic DNA sequencing of MDCK-II wild-type (WT) and ARHGAP12 knockout (KO) lines. The guide RNA (indicated in blue) targets exon 3. (B) WB analysis of cell lysates from MDCK-II WT and ARHGAP12 KO lines. Clones F9, F21, and F23 show a complete loss of ARHGAP12 expression. No changes in protein expression were detected for the proteins tested. (C) Immunofluorescence analysis of MDCK-II ARHGAP12 KO lines F9 and F21 co-cultured on glass with MDCK-II WT cells. Cells were fixed and stained with antibodies against ARHGAP12 and ZO-1. KO cells were identified via the GFP signal. Maximum intensity projections of confocal z-stacks are shown. Note that nuclear ARHGAP12 signals are due to non-specific ARHGAP12 antibody reactivity. (D) MDCK-II WT and ARHGAP12 KO F21 cells were co-cultured on Transwell filters for 10 days. Cells were fixed and stained with anti-ZO-1 (top panel) or anti-Occludin (bottom panel) antibodies. Maximum intensity projections of confocal z-stacks are shown.

**Figure S8:**
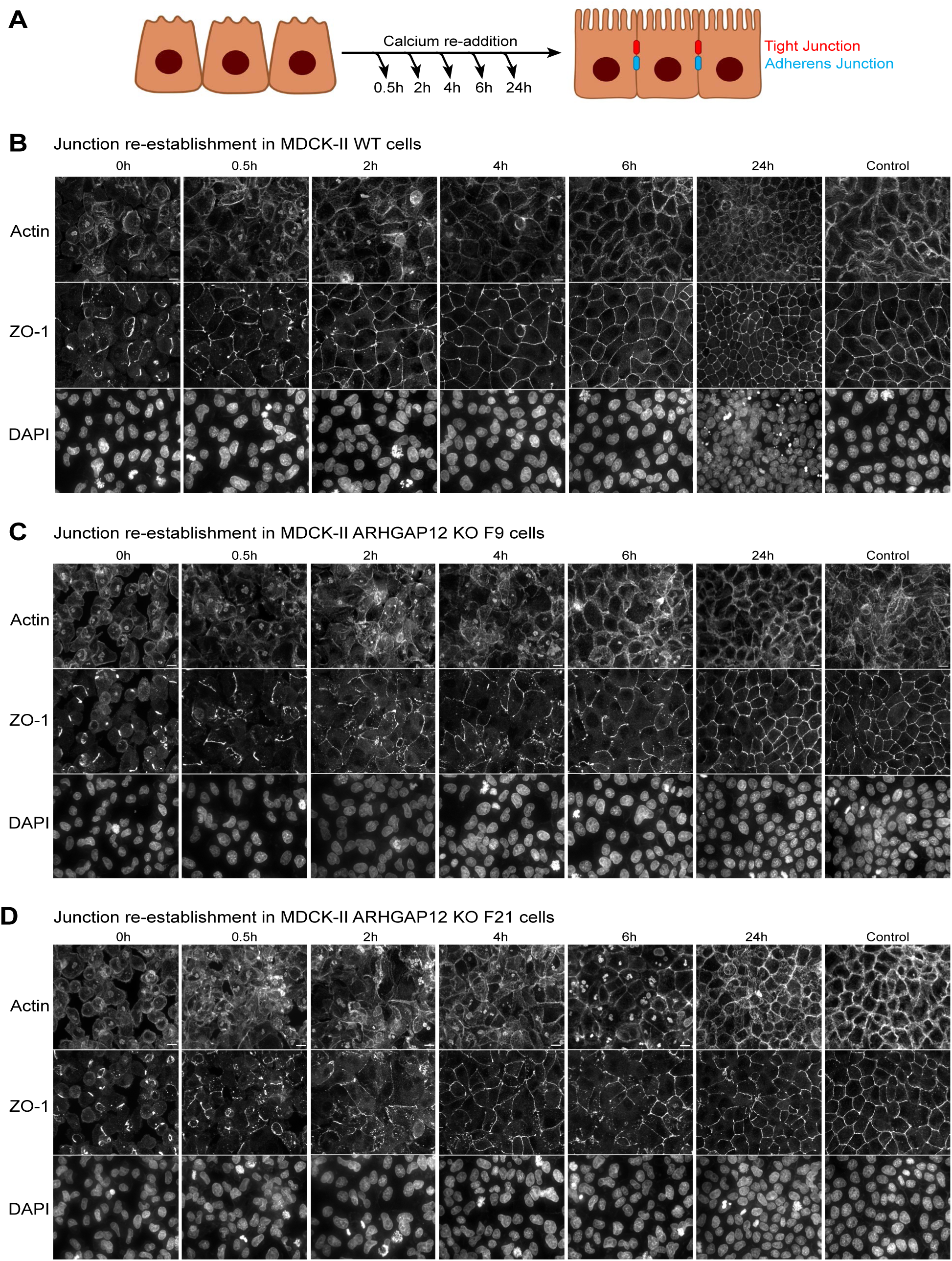
ARHGAP12 promotes TJ formation. (A) Schematic illustrating the calcium switch assay. Cell monolaters are depleted of Ca^++^ for 16-24 hrs, resulting in the dissociation of cell junctions. Re-addition of Ca^++^ permits cell junctions to reform over time. (B-D) Calcium switch assay of MDCK-II WT (B), ARHGAP12 KO F9 (C), and ARHGAP12 KO F21 cells (D) grown on coverslips. Cells were fixed at different time points (0, 0.5, 2, 4, 6, and 24 hrs) and stained with Phalloidin to visualize actin and antibodies against ZO-1 to visualize TJ. DAPI was used to visualize nuclei. Scale bars 10 µm.

## Notes

### Competing Interest Statement

The authors have declared no competing interest.

